# Context Dependence of Biological Circuits

**DOI:** 10.1101/360040

**Authors:** Thomas A. Catanach, Reed McCardell, Ania-Ariadna Baetica, Richard M. Murray

## Abstract

It has been an ongoing scientific debate whether biological parameters are conserved across experimental setups with different media, pH values, and other experimental conditions. Our work explores this question using Bayesian probability as a rigorous framework to assess the biological context of parameters in a model of the cell growth controller in You *et al*. When this growth controller is uninduced, the *E. coli* cell population grows to carrying capacity; however, when the circuit is induced, the cell population growth is regulated to remain well below carrying capacity. This growth control controller regulates the *E. coli* cell population by cell–cell communication using the signaling molecule AHL and by cell death using the bacterial toxin CcdB.

To evaluate the context dependence of parameters such as the cell growth rate, the carrying capacity, the AHL degradation rate, the leakiness of AHL, the leakiness of toxin CcdB, and the IPTG induction factor, we collect experimental data from the growth control circuit in two different media, at two different pH values, and with several induction levels. We define a set of possible context-dependencies that describe how these parameters may differ with the experimental conditions and we develop mathematical models of the growth controller across the different experimental contexts. We then determine whether these parameters are shared across experimental contexts or whether they are context-dependent. For each of these possible context-dependencies, we use Bayesian inference to assess its plausibility and to estimate the growth controller’s parameters assuming this context-dependency. Ultimately, we find that there is significant experimental context-dependence in this circuit. Moreover, we also find that the estimated parameter values are sensitive to our assumption of a context relationship.

## 1 Introduction

The goal of synthetic biology has been to engineer reliable circuits composed of standardized parts that can easily be combined together [1, 7, 16, 18]. However, it has been discovered that context dependence in synthetic biological circuits causes parts and modules to behave unpredictably in the cell under different experimental conditions, as discussed in [12, 23, 24]. Differences in design, implementation in a different model organism [17], or different experimental conditions can all have unexpected results in the functioning of synthetic circuits [15]. Poorly understood context dependence of biological parameters limits synthetic biologists’ ability to construct reliable, large-scale circuits [20]. We investigate the context dependence of the biological parameters of the cell growth controller in You *et al*. [25] using Bayesian methods.

We define context dependence to mean that the system is dependent on some condition or context in a way that is not explicitly mathematically modeled. This could be because the relationship is very complicated and challenging to model or because we do not aim to model it in that much detail. Thus, context dependence represents an unmodeled relationship. We can learn the effects of context on our parameters even without developing a model of it by understanding the variability in model parameters between different contexts. For example, in this paper we consider context relationships with respect to different control circuit implementations, different pH values, and different levels of induction. The biological parameters of this circuit have complicated dependencies on these variables that would be challenging to model in detail. Constructing mathematical models to characterize these dependencies goes beyond our current knowledge. However, using Bayesian system identification methods, we are able to tease out relationships of context dependence and parameter variability that can provide valuable information for future experiments [3, 14].

Identifying the biological circuit’s context can also help develop better models of a new circuit based on previously available experimental data and on models of similar circuits. Typically, to model a new circuit, we use models and parameter values that were measured for similar circuits under different experimental conditions. Parameter estimates may sometimes be improved using system identifcation methods to infer values for the new circuit from experiments under different conditions or different control implementations [19, 21]. This combination of results from different experimental conditions and different circuits leads to many questions about the role of context dependence. If there is context dependence, combining these results without considering the role of context can cause significant bias. This means that the models that we learn through system identification poorly capture the true structure of the biological circuit and poorly predict its behavior under different conditions.

In this paper, we study whether biological parameters are conserved across experimental conditions with different pHs, IPTG induction levels, and control experiment implementations of the cell growth controller in You *et al*. [25]. This circuit controls cell growth using cell–cell communication and toxin-mediated cell death [10, 11, 13]. The signaling molecule AHL (Lux) senses the cell density and the circuit responds to high cell density by killing cells using toxin CcdB. When induced with isopropyl–d-thiogalactopyranoside (IPTG), plasmid pLac-LuxR-LuxI expresses regulatory proteins LuxI and LuxR. The controller responds to activated LuxR by expressing toxin CcdB to induce cell death. We investigate the context dependence of parameters such as the cell growth rate, carrying capacity, AHL degradation rate, leakiness of AHL, and leakiness of toxin CcdB on experimental conditions using mathematical models of the circuit.

We define mathematical models that relate the dynamics of the cell population to biological parameters of the circuit and also a set of possible context relationships that determine which parameters are sensitive to changes in experimental conditions. The significance of these context relationships is captured using parameters that define how much the experimental condition affect the biological parameters’ value. Therefore, when learning the model of the circuit, we must identify both the biological parameters that describe the dynamics of the circuit and parameters that describe the context variability. Bayesian parameter inference uses the experimental data to identify the model parameters, while Bayesian model selection compares the different possible context relationships [3, 6, 14]. Bayesian model selection has been used in many areas, including systems biology, to compare different candidate mathematical models, but not to investigate context dependence [2, 5, 6, 22].

To assess whether these biological parameters are conserved, we collect experimental data from the You *et al*. circuit in two different growth media, Lysogeny Broth (LB) and Tryptone Buferred KCl (TBK). Further, motivated by the pH dependence observed in [25], we run the circuit at two pH values of 6.6 and 7.4 and also at different IPTG induction levels. Additionally, we perform two control experiments with an incomplete circuit that contains only one of the two plasmids pLux-CcdB and pLac-LuxR-LuxI. We use fluorescence measurements to quantify the cell population over time.

Using the Bayesian framework, we are able to quantify context dependence, to identify parameters that vary between implementations, and to assess the sensitivity of the parameter estimates to different context models based on the experimental data. We identify context relationships and estimate the model parameters with a variant of Markov Chain Monte Carlo called Sequential Tempered Markov Chain Monte Carlo [8]. We find that for the different control experiments in the LB medium, there is significant context dependence of the growth rate and of the carrying capacity parameters, which agrees with our intuition about the biological system. Further, in the TBK medium, we identify significant context dependence among the biological parameters with respect to both changes in pH and in IPTG induction. However, not only do we find significant context dependence, but we also observe that our assumptions about the context relationship can significantly influence the parameters values we find using Bayesian parameter inference from the experimental data. The sensitivity of the posterior parameter distribution to our assumed model of context highlights how critical it is to include context uncertainty into the model used to identify parameters from the data. Therefore, the primary contribution of this work is to show that Bayesian statistics provides a powerful framework for quantitatively considering context dependent relationships and that it can help inform future experiments to improve models of biological systems.

This paper is organized as follows: Section 2 describes the biological circuit, its experimental implementation and its mathematical model. Section 3 describes Bayesian methods and how they can be used to infer the biological parameters’ context. Section 4 describes the results of inferring the context. Section 5 provides details about materials and methods. Finally, Section 6 concludes. The supplemental material contains information about the numerical results and about Bayesian probability theory.

## 2 The Cell Growth Regulation Circuit

To examine the question of the context dependence of biological parameters, we first describe the growth regulation circuit and model its dynamics across multiple experimental setups. Then, in each experimental setup, we collect fluorescence data measurements for a population of cells that carry the growth regulation circuit. Finally, we combine the mathematical models and the data measurements to infer the biological parameters’ context dependence as discussed in Section 3.

### 2.1 The description of the cell growth regulation circuit

**Figure 1:**
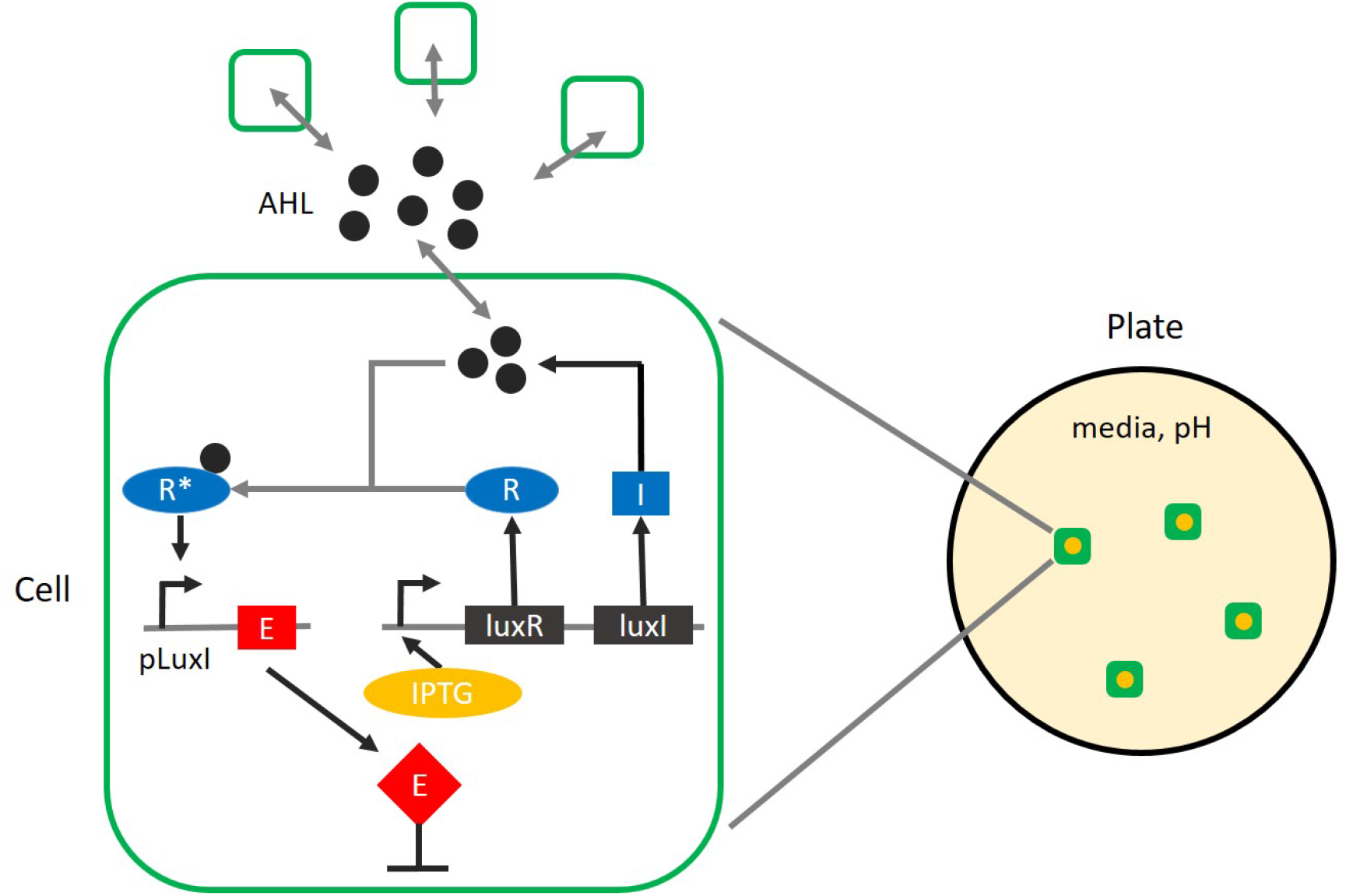
A schematic of the circuit for cell growth control. The cell population is controlled using cell-cell communication with quorum sensing proteins and toxin-mediated cell death. The toxin gene is denoted by E and it expresses the CcdB to the cells (red). Filled black circles represent quorum sensing proteins AHL. The notation I, R and R* represents proteins LuxI, LuxR and active LuxR, respectively (blue). Filled orange circles represent molecules of the inducer IPTG that activates the circuit by turning on LuxI and LuxR. The growth circuit is implemented with two plasmids, pLac-LuxR-LuxI and pLuxCcdB, with resistance to antibiotics Chloramphenicol and Kanamycin, respectively [25]. They are cloned into the *E. coli* cell and the cell expresses the growth circuit only when the two plasmids are simultaneously present. The cell culture is grown on a plate with one of two media (LB or TBK) at one of four different IPTG levels. Additionally, the TBK medium uses two pH concentration levels of 6.6 and 7.4.

### 2.2 Data measurement description

We grow *E. coli* cells with both the pLux-ccdB and the pLac-LuxR-LuxI plasmids from [25] in two media (LB and TBK) at four IPTG induction levels (0mM, 0.2mM, 1mM, 5mM). The experiments with no IPTG induction (IPTG = 0mM) are control experiments in which the circuit should not function. In the TBK medium, we also grow cells at two pH levels of 6.6 and 7.4 to test context dependence on pH. In the LB medium, we additionally run two control experiments in which only one of the plasmids is present in the cells. We collect fluorescence measurement data every 7 minutes for 21 hours in the LB medium and for 28 hours in the TBK medium. The data measurements are illustrated in Figures 2 and 3. For more details on the experimental method, we direct the reader to Section 5.

**Figure 2:**
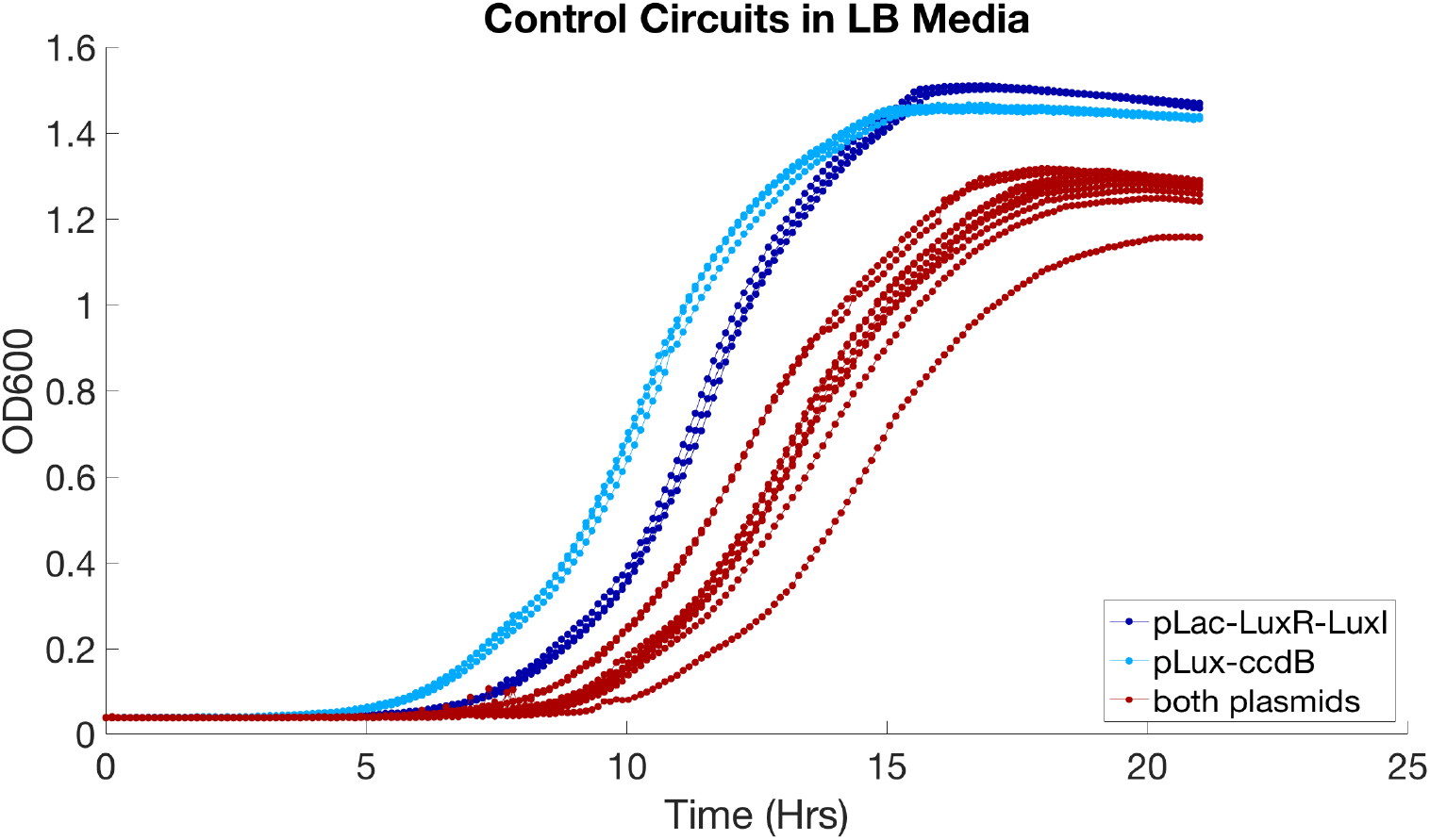
Fluorescence data measurements for the control experiments of the growth circuit in the LB medium. OD600 absorbance measurements for multiple experimental setups of the growth control circuit. Two control circuits are only partially implemented circuits with only one of the two plasmids present, pLac-LuxR-LuxI or pLux-CcdB. The other control circuit is the full circuit with both plasmids present, but is uninduced. Lines of the same color indicates replicate experiments run under the same experimental conditions.

**Figure 3:**
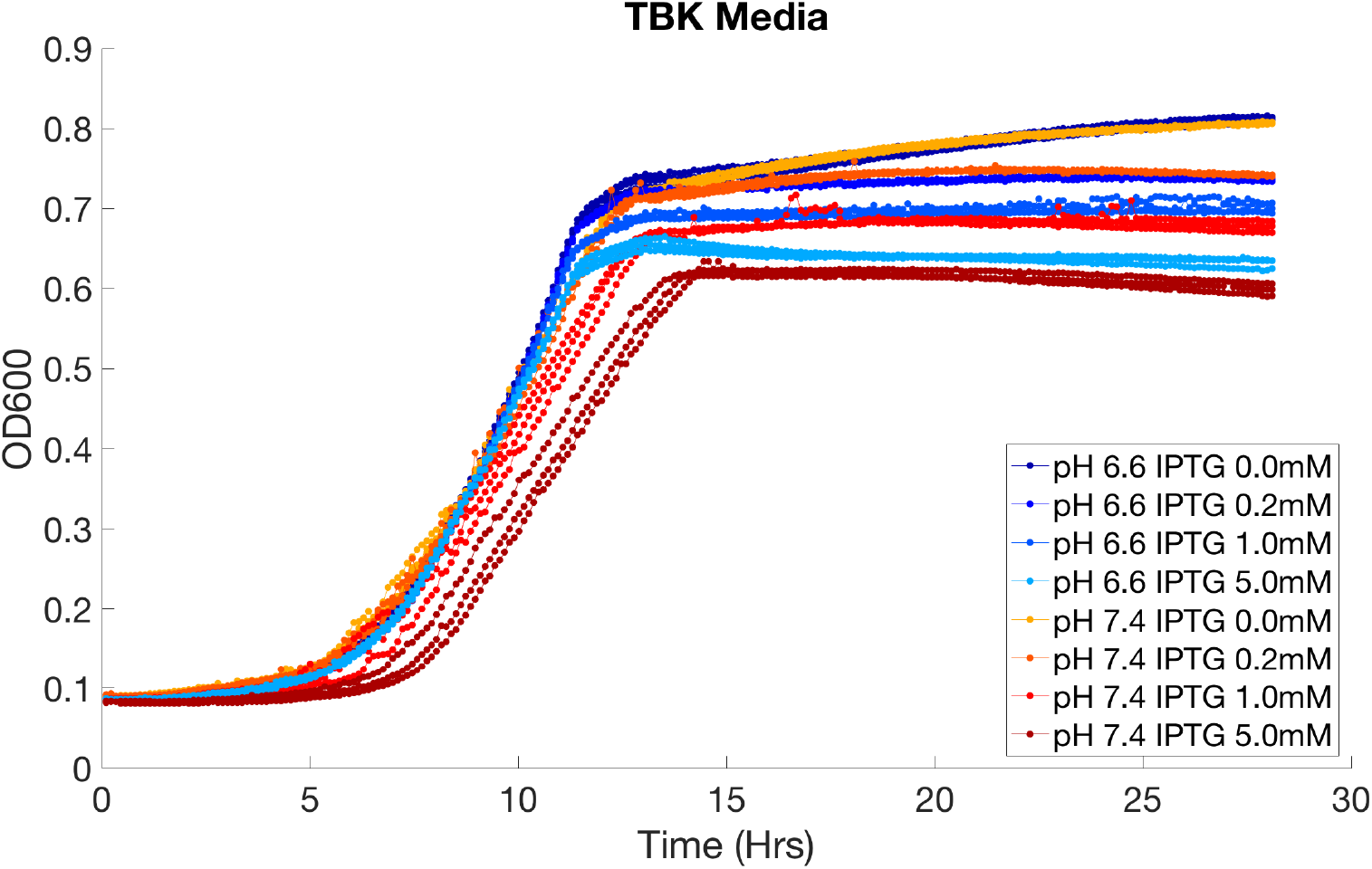
Fluorescence data measurements for the growth circuit in the TBK medium. OD600 absorbance measurements of the growth circuit at four different IPTG induction levels of 0mM, 0.2mM, 1.0mM, and 5.0mM and at two pH levels of 6.6 and 7.4. The circuit runs for 28 hours in the TBK medium. Lines of the same color indicates replicate experiments run under the same experimental conditions.

### 2.3 Mathematical models of the growth circuit

We develop mathematical models for the different circuits that we collect experimental data for. We describe models of the three control experiments, which include two incomplete versions of the full circuit and the uninduced (IPTG = 0mM), but complete circuit. Subsequently, we describe the model of the complete and induced circuit. These models are able to capture the dynamics of the cell population in a variety of media, pH, and IPTG induction levels by adjusting the parameter constants in a modified logistic equation system. The parameter constants in the equations used to model these circuits are described in Table 1.

**Table 1:** The list of species and reaction rates in the growth circuit model. We consider the following species in our modeling: the cell density (N), the signaling molecule AHL (A), and the toxin CcdB (E). We model cell growth (rate k) and death (d), the signaling molecule production (*v_A_*), degradation (*d_A_*), and induction (Hill coefficient n), as well as toxin production (*k_E_*) and degradation (*d_E_*). We also include the leakiness of the toxin (*t*_0_) and of the signaling molecule (*A*_0_).

| Chemical Species | Species Notation | Chemical Rates | Rates Notation |
| --- | --- | --- | --- |
| cell number | $N$ | growth rate | $k$ |
| toxin | $E$ | death rate | $d$ |
| AHL | $A$ | toxin induction | $k_E$ |
| IPTG | $I$ | toxin degradation | $d_E$ |
| basal toxin | $t_0$ | AHL induction | $v_A$ |
| basal AHL | $A_0$ | AHL degradation | $d_A$ |
| carrying capacity | $N_m$ | Hill coefficient | $n$ |

First, we model the control experiments where only one plasmid is present in the cells. Since the growth circuit is uninduced and inactive, the cell population number in the two control experiments is modeled by the logistic equation

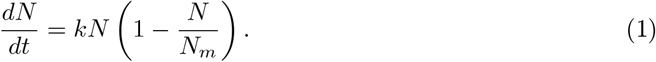

Secondly, we extend the model in [25] to capture the experiments when the growth regulation circuit is induced. We model cell growth and death, the signaling molecule production, degradation, and induction, as well as toxin production and degradation. We assume that the cell density follows the logistic equation, but modified to reflect the cells dying in proportion to toxin concentration. We assume that there is leakiness of both the signaling molecule and the toxin (represented by *A*_0_ and *t*_0_). Additionally, we assume that the IPTG induction affects the concentration of AHL in the form of a Hill function and that it is proportional to the cell density. Furthermore, we assume that the cooperativity of the IPTG interaction is one. The production rate of toxin is assumed to be proportional to AHL concentration. We refer the reader to [25] for other assumptions underlying the modeling. The equations of the induced growth regulation circuit are as follows:

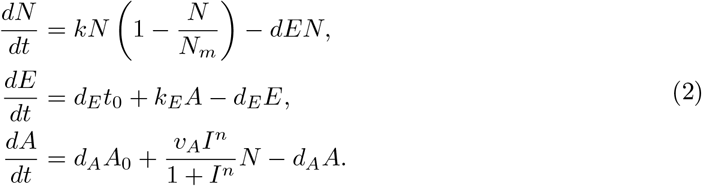

Lastly, the model for the LB and the TBK media with no IPTG induction is a simplification of Equation (2), where we set the species *I* to value zero:

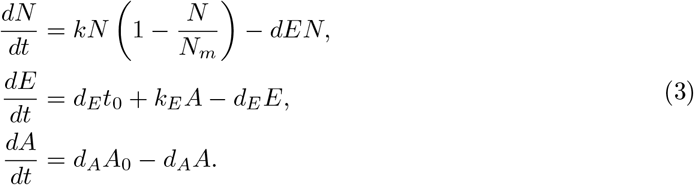

For these models, we need to estimate the value of many of the parameters and to identify how these parameters change with the implementation context in the experiments. Therefore, we use the data measurements in Section 2.2 and the three models in Section 2.3 to determine the biological context of the circuit’s parameters using the methods of Bayesian inference and model selection.

## 3. Bayesian Inference And Context Models

Within this section, we investigate how biological context affects the parameters in the circuit models from Section 2.3 without deriving a mathematical model that explicitly defines the relationship between the implementation contexts and the biological parameters. In order to learn these un-modeled relationships, we first need to describe a set of possible context dependence models that define which parameters are sensitive to each context and to specify the strength of the sensitivity. Then we can identify which context relationship model best describes the measurement data from the experiments in Section 2.2 using the Bayesian framework. This approach can be broken up into two steps. First, we assume a context dependence relationship and we find parameter values in our model that agree with the data. Second, for all the context models, we consider which ones match the data without over-fitting. The first problem can be solved using Bayesian parameter estimation and the second can be solved using Bayesian model selection. These topics have been extensively covered in [3, 14]. However, their application to the inference of context dependence in biological systems is novel.

### 3.1 Bayesian inference

The process of Bayesian inference for parameter estimation is visualized in Figure 4. Using Bayesian inference, we find plausible parameter values for a mathematical model that agree with the data and with our prior knowledge. These notions of plausibility and agreement are rigorously defined and quantitative using the language of probability and probability distributions. Ultimately, we construct a probability distribution, called the posterior distribution, over model parameters that can then be used to estimate properties of the system. Parameters with high probability according to this distribution are those that fit the data well.

**Figure 4:**
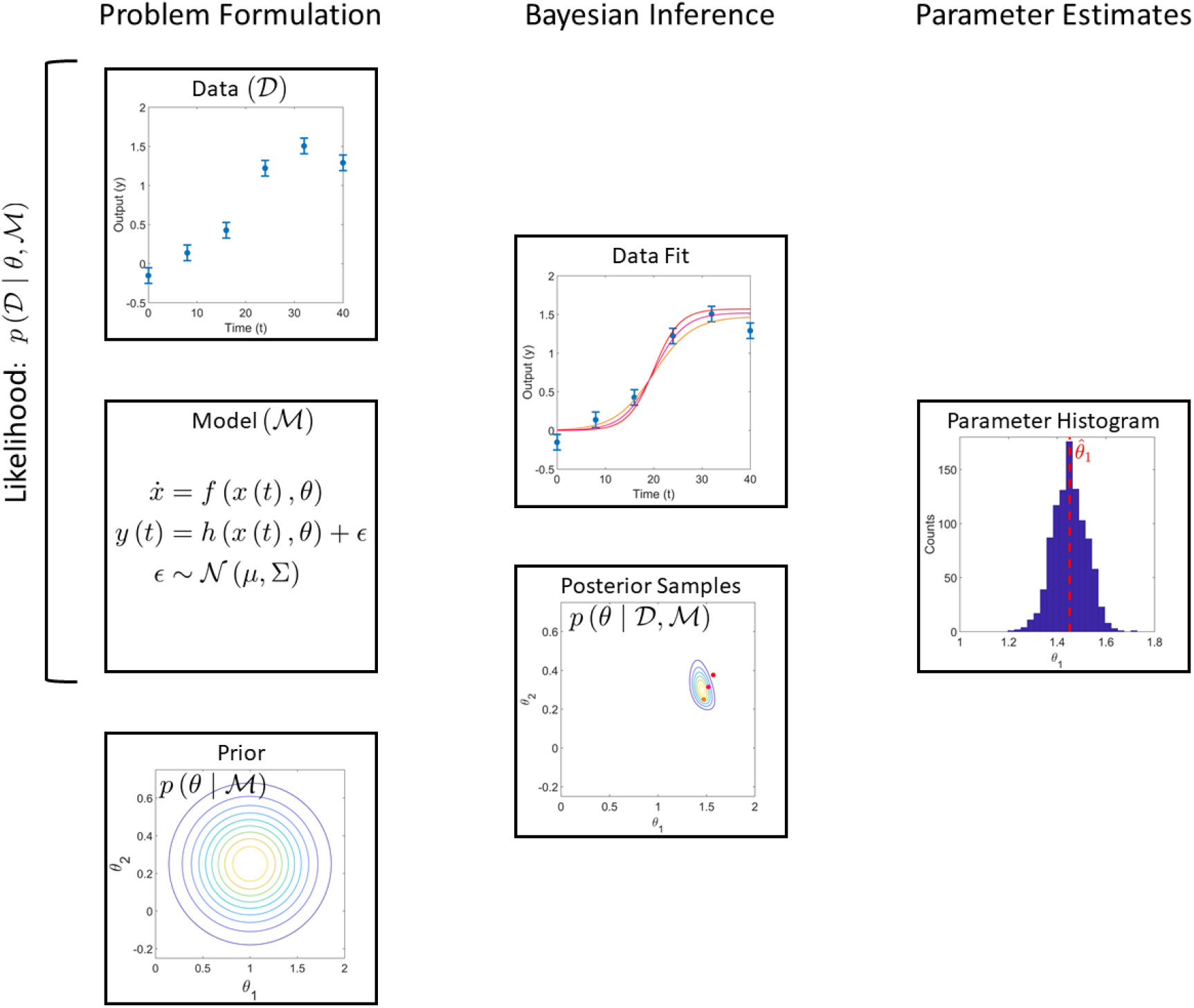
A schematic of Bayesian parameter estimation. The goal of Bayesian parameter estimation is to find a probability distribution that describes the range of model parameter values, *θ*, that fit the experimental data, 𝒟. The formulation begins with a prior probability distribution for the parameter values, *p* (*θ* | ℳ), a mathematical model of the system, ℳ, and the set of data measurements from experiments. The prior distribution captures our initial guess of which values are more likely than others for a model parameter. This probability distribution will be updated using a function which combines the model and the experimental data to get better estimates of which parameter values are more likely according to the data. This function, known as the likelihood, *p* (𝒟 | *θ*, ℳ), is higher for parameters that cause the model to predict and fit the data better. The updated probability distribution is known as the posterior probability distribution, *p* (*θ* | 𝒟, ℳ). Typically, we cannot work directly with the posterior distribution, but we can work with samples from the distribution that represent likely parameter values which yield good fits to the data. From the posterior sample population, we can estimate properties of the parameter’s posterior distribution such as its mean or standard deviation.

In order to construct the posterior probability distribution, *p* (*θ* | 𝒟, ℳ), over model parameters, *θ*, for a mathematical model ℳ and experimental data 𝒟, we define a prior probability distribution, *p* (*θ* | 𝒟, ℳ), and then update this distribution based on the data. The prior distribution defines our initial assumptions about the model parameters. The prior distribution is updated to construct the posterior distribution using Bayes’ Theorem to improve our estimate of the model parameter vector by conditioning the distribution on the observational data 𝒟 according to the equation:

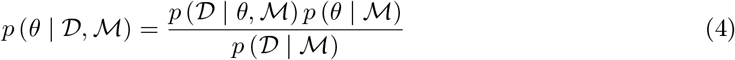

The updating step is performed through the likelihood function, *p* (𝒟 | *θ*, ℳ), which assesses the fit of the data given the model ℳ and its parameters *θ*. Therefore, the highest probability parameters of the posterior distribution are those which both fit the data well and have a high probability according to the prior. The normalization factor *p* (𝒟 | ℳ), which is necessary to ensure this is a valid probability distribution, also provides information about the likelihood of the model, ℳ, which can then be used to compare different mathematical and context dependence models. Therefore, it is also known as the model evidence.

In most practical implementations of Bayesian inference algorithms, we cannot analytically manipulate these probability distributions directly but can only use samples from these distributions. However, using these samples, we can estimate properties of the model such as the mean and standard deviation of a model parameter and we can make predictions about the behavior of the modeled system under different conditions. For more details about formulating and solving Bayesian inference and model selection problems, see Appendix B.

### 3.2 Inferring models of context

Using probabilistic models, we can capture the uncertainty introduced by complex relationships between biological systems and their implementation context. These relationships are too complicated to model explicitly; however, simpler probabilistic models of these context relationships allow us to capture the uncertainty that comes form not modeling the relationships directly. In these probabilistic context models, if a parameter from the mathematical model of the biological system is context dependent, then its value varies according to the probability distribution when the context of the biological system changes. The more uncertain the value of the parameter, the more significant the context dependence. In contrast, a context independent parameter has a fixed value as the context changes and therefore its value is shared between all mathematical models of experimental contexts. Learning about the context dependence of a system involves deciding which parameters are context dependent and assessing the strength of the dependence by learning the parameters using the prior. This formulation is visualized using the graphical models in Figure 5.

**Figure 5:**
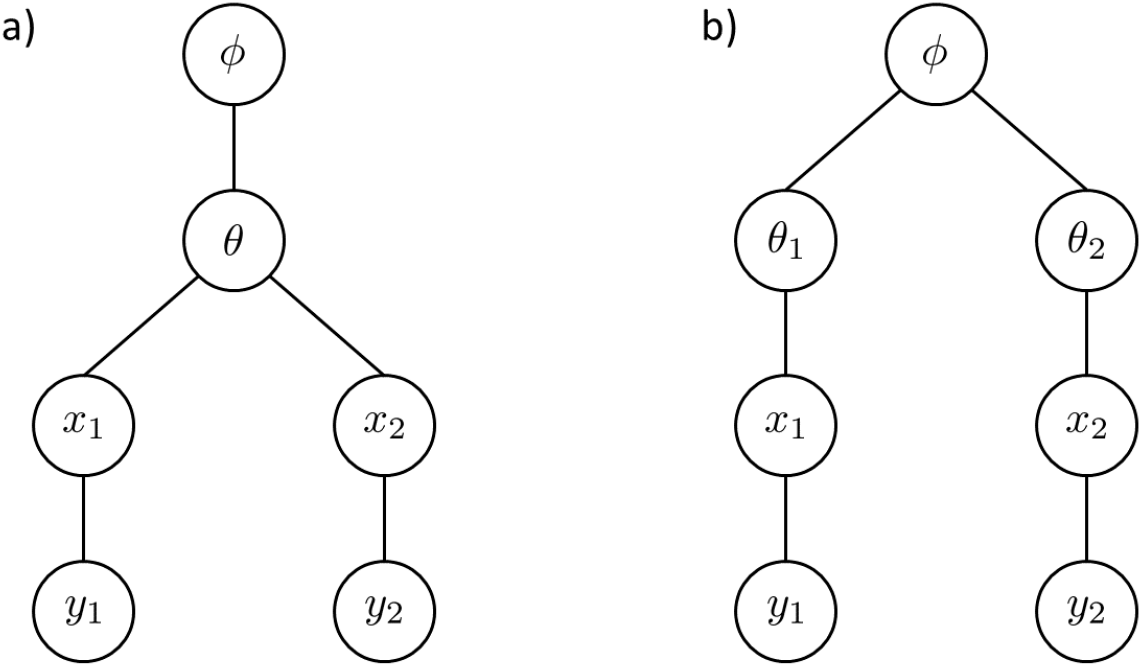
A comparison of two experimental context relationships. Graphical models can be used to express probabilistic relationships between variables such as context dependence. The lines between variables represent probabilistic dependencies. In this example, *y*_1_ and *y*_2_ denote measurements from two different experiments under different conditions such as different pH or inducer values. *x*_1_ and *x*_2_ are the state variables for each experimental setup (i.e. the numbers of cells). *θ*, *θ*_1_, and *θ*_2_ are parameters in the mathematical model of the system that describe the evolution of the state variables, such as the cell growth rate or the carrying capacity in the logistic growth model. *ϕ* are parameters that describe the prior distribution for the variables *θ* such as their prior mean and standard deviation. The graphical models a) and b) represent two different context relationships between the two experimental conditions by modeling different probabilistic relationships between the variables. In model a), the parameter *θ* is shared between the two experimental conditions and thus expresses independence of the context. In model b), parameters *θ*_1_ and *θ*_2_ are different between the models for the two experiments and therefore express context dependence. *θ*_1_ and *θ*_2_ are still linked through the common parameters *ϕ* that can capture the strength and nature of the context dependence.

First, to infer the appropriate context dependence for each parameter, we hypothesize a set of models, each with a different context relationship. After we perform Bayesian parameter estimation to identify parameters using several different candidate context dependency models, the next step is to judge which context model is most appropriate, a problem known as Bayesian model selection. Different context relationship models are evaluated based on how closely they fit the data and on how much information was needed to update their prior distributions to get the posterior distribution. The trade-off between these two objectives is called the Bayesian Occam’s Razor and it is a rigorous probabilistic method to express over-fitting [3]. For each context relationship model, we compute its model evidence and we compare them to assess the relative likelihoods of the different context models. Appendix B discusses these probabilistic models and model selection in more detail.

We use these two steps to learn about the biological context of parameters in the mathematical models used to describe the biological circuit in [25]. The parameters in the model from Section 2.3 are either shared or dependent on context variables such as the pH or IPTG induction levels. For the different context dependency relationships, the parameters are inferred using Bayesian inference based on the data collected from experimental runs for different contexts. Then, for each context dependence model, the relative plausibility is computed to find which best represents the parameter context relationships.

## 4 Results Of The Method

### 4.1 Determining context dependence for the control experiments in the LB medium

In this subsection, we determine whether the cell growth rate and the cell population’s carrying capacity are context-dependent in three control experiments in the LB medium. The three control experiments we consider are as follows: two control experiments where we run the growth control circuit with only one of the two plasmids present, and an additional experiment where we run the circuit with both plasmids present but with no IPTG induction. In all three control experiments, the growth control circuit should not be functional. More details on the control experiments were provided in Section 2.2. Equation (1) in Section 2.3 models the behavior of the circuit with only one plasmid since the growth control is not fully implemented. Equation (3) models the uninduced circuit since the growth control is implemented without IPTG induction. The growth rate and the carrying capacity are the only parameters common between these two mathematical models.

If a biological parameter in the mathematical models in Equations (1) and (3) is context dependent on the experimental conditions, then the model for each of these three control experiments has a different value for the parameter. The values for this parameter across each circuit are linked by a shared hyperprior distribution that controls the significance of the context dependence. We consider three types of context dependence: no dependence, growth rate dependence, and growth rate and carrying capacity dependence, as described in Table 2. For each of the three context relationship models, the prior parameter distributions are the same. The posterior parameter distributions are then sampled using MCMC to find plausible parameters that fit the data. For these three context relationship models, we also compute the relative model probabilities, as illustrated in Table 2. We obtain that both the growth rate and the carrying capacity are context dependent. Samples from the distribution of the inferred growth rate and carrying capacity values are plotted in Figure 6. This figure shows significant separation in parameter space between the inferred parameters for the three control experiments, which is indicative of context dependence. This result is realistic from a biological perspective since having only one of the two plasmids that implement the growth control circuit would affect the cell growth rate and the cell population’s carrying capacity.

**Figure 6:**
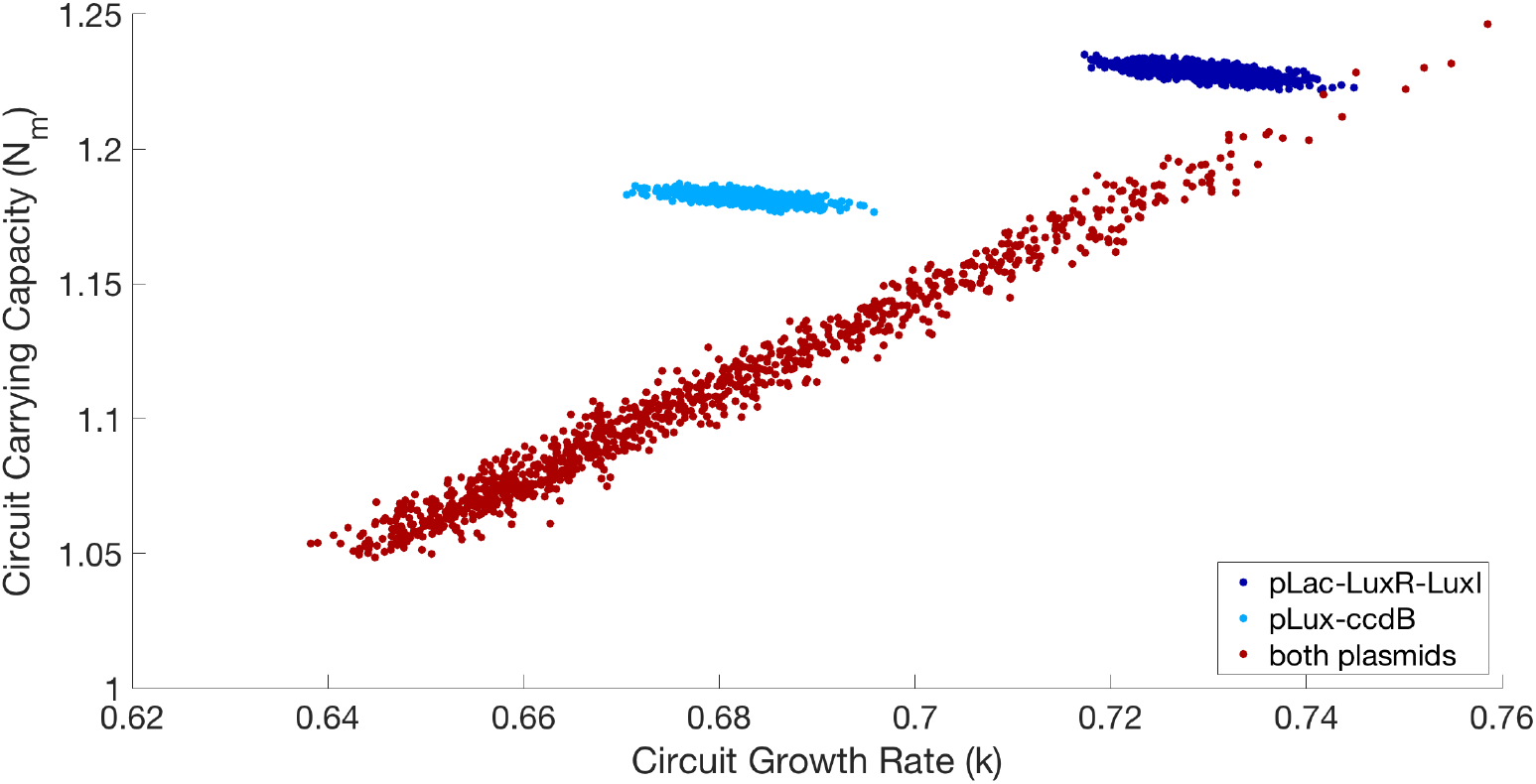
Posterior samples of the cell growth rate and carrying capacity parameters for the three control experiments in the LB medium. The two control experiments with only one plasmid present, pLac-LuxR-LuxI or pLux-CcdB, have significantly different growth rates and carrying capacities. The control experiment with both plasmids present but no IPTG induction has a posterior which is less informed by the data due to the added complexity of the model; however, the bulk of its probability mass is in a region of parameter space far from the other two control experiments. This is indicative of significant context dependence of the growth rate and the carrying capacity for these three control experiments.

**Table 2:** Models of context for the control experiments in the LB medium. "Shared" parameters are the same between the control experiment contexts, while "Dependent" parameters are different. The probability of a model is the relative probability based on Bayesian model selection. We find that the cell growth rate and the cell population’s carrying capacity are context dependent, with probability close to one.

| Model | Growth Rate | Carrying Capacity | Basal Toxin | Probability |
| --- | --- | --- | --- | --- |
| Model 1 | Shared | Shared | Shared | $10^{-191}$ |
| Model 2 | Dependent | Shared | Shared | $10^{-66}$ |
| Model 3 | Dependent | Dependent | Shared | $\approx 1.0$ |

### 4.2 Determining context dependence on IPTG induction and pH values in the TBK medium

We next investigate context dependence in the growth control circuit with respect to pH values and to IPTG induction levels. We are motivated by You *et al*.’s finding that pH affects the circuit’s performance and may lead to context dependent parameters [25]. We infer parameters for a set of eleven context relationships models described in Table 3 and we find their relative probabilities. Each parameter can have one of four possible context relationships: a shared value between all contexts i.e. no context relationship, pH context dependence, IPTG context dependence, or both pH and IPTG dependence. For all context relationship models, the Bayesian problems share the same hyperprior distribution for a parameter’s prior to make reasonable comparisons. The structure of the prior and hyperprior distribution are found in Tables 4 and 5 in the Appendix.

**Table 3:** Models of context for the dependence on pH level and IPTG induction level. "Shared" parameters are the same regardless of the pH and the IPTG values, while parameters labeled with "pH" and/or "IPTG" vary with that particular context between experimental conditions. The probability of a model is the relative probability based on Bayesian model selection. We find that the most probable model is Model 11, which describes the context dependence of the cell growth rate, the cell population carrying capacity, and the AHL induction on both the pH level and the IPTG induction value.

| Model | $k$ | $N_m$ | $v_A$ | $d_A$ | $t_0$ | $A_0$ | Probability |
| --- | --- | --- | --- | --- | --- | --- | --- |
| Model 1 | Shared | Shared | Shared | pH | Shared | Shared | $10^{-241}$ |
| Model 2 | pH | Shared | Shared | pH | Shared | Shared | $10^{-239}$ |
| Model 3 | pH | pH | Shared | pH | Shared | Shared | $10^{-230}$ |
| Model 4 | Shared | Shared | pH IPTG | pH | Shared | Shared | $10^{-207}$ |
| Model 5 | pH | Shared | pH IPTG | pH | Shared | Shared | $10^{-204}$ |
| Model 6 | pH | pH | pH IPTG | Shared | Shared | Shared | $10^{-202}$ |
| Model 7 | pH | pH | pH IPTG | pH | Shared | Shared | $10^{-195}$ |
| Model 8 | pH | pH | IPTG | pH | Shared | Shared | $10^{-204}$ |
| Model 9 | pH | pH | pH IPTG | pH | pH | pH | $10^{-196}$ |
| Model 10 | pH IPTG | pH | pH IPTG | pH | Shared | Shared | $10^{-58}$ |
| Model 11 | pH IPTG | pH IPTG | pH IPTG | pH | Shared | Shared | $\approx 1.0$ |

**Table 4:** The prior distributions of biological parameters. The priors for the biological parameters of the model in Section 2.3. The priors for *k*, *N_m_*, *v_A_*, *d_A_*, *t*_0_, and *A*_0_ are defined using hyperparameters and are influence by the choice of context model. The priors for *N* (0), *E* (0), *A* (0), and *σ* are fixed and not influenced by the choice of context model.

| Parameter | Type | Description |
| --- | --- | --- |
| $k$ | Log-normal | Lognormal ($\mu_k, \sigma_k$) |
| $N_m$ | Uniform | Unif [$a_{N_m}, a_{N_m} + r_{N_M}$] |
| $v_A$ | Log-normal | Lognormal ($\mu_{v_A}, \sigma_{v_A}$) |
| $d_A$ | Log-normal | Lognormal ($\mu_{d_A}, \sigma_{d_A}$) |
| $t_0$ | Exponential | Exponential ($\lambda_{t_0}$) |
| $A_0$ | Exponential | Exponential ($\lambda_{A_0}$) |
| $N(0)$ | Uniform | Unif [0.1, 0.2] |
| $E(0)$ | Exponential | Exponential (0.2) |
| $A(0)$ | Exponential | Exponential (0.2) |
| $\sigma$ | Log-normal | Lognormal (17, 0.5) |

**Table 5.**
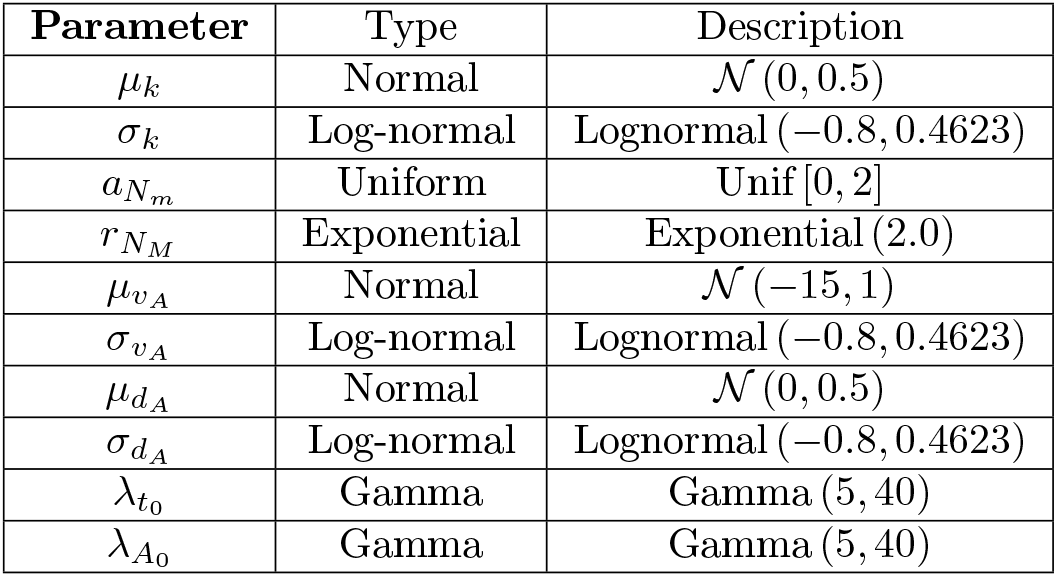
Prior distribution of the hyperparameters. The hyperparameters define the prior distribution (Table 4) of some of the biological parameters in Section 2.3. The hyperparameter priors define our current beliefs about the ranges of the biological parameter values and the variability of the parameter values introduced by context dependence.

| Parameter | Type | Description |
| --- | --- | --- |
| $\mu_k$ | Normal | $\mathcal{N}(0, 0.5)$ |
| $\sigma_k$ | Log-normal | Lognormal $(-0.8, 0.4623)$ |
| $a_{N_m}$ | Uniform | Unif $[0, 2]$ |
| $r_{N_M}$ | Exponential | Exponential (2.0) |
| $\mu_{v_A}$ | Normal | $\mathcal{N}(-15, 1)$ |
| $\sigma_{v_A}$ | Log-normal | Lognormal $(-0.8, 0.4623)$ |
| $\mu_{d_A}$ | Normal | $\mathcal{N}(0, 0.5)$ |
| $\sigma_{d_A}$ | Log-normal | Lognormal $(-0.8, 0.4623)$ |
| $\lambda_{t_0}$ | Gamma | Gamma (5, 40) |
| $\lambda_{A_0}$ | Gamma | Gamma (5, 40) |

For each of the possible context relationship models considered, we sample the posterior distribution from Bayesian parameter inference to find plausible parameter values. Based on the posterior parameter distributions from the different model contexts, we can draw conclusions about different parameter values. The expected values of the posterior for the growth rate, the carrying capacity, and the AHL induction factor for different context models are shown in Figures 7 – 9. We observe that some parameters, such as the growth rate, change significantly depending on the context model, while other parameters, such as the carrying capacity or the AHL induction rate, are fairly consistent, indicating that their values are inferred reliably. This result highlights the importance of considering context when performing Bayesian inference for biological parameters since the inferred value of some parameters may be quite sensitive to context. The histograms corresponding to the posterior distributions of the biological parameters with different context models are shown in Figures 16 – 18 in the Appendix.

**Figure 7:**
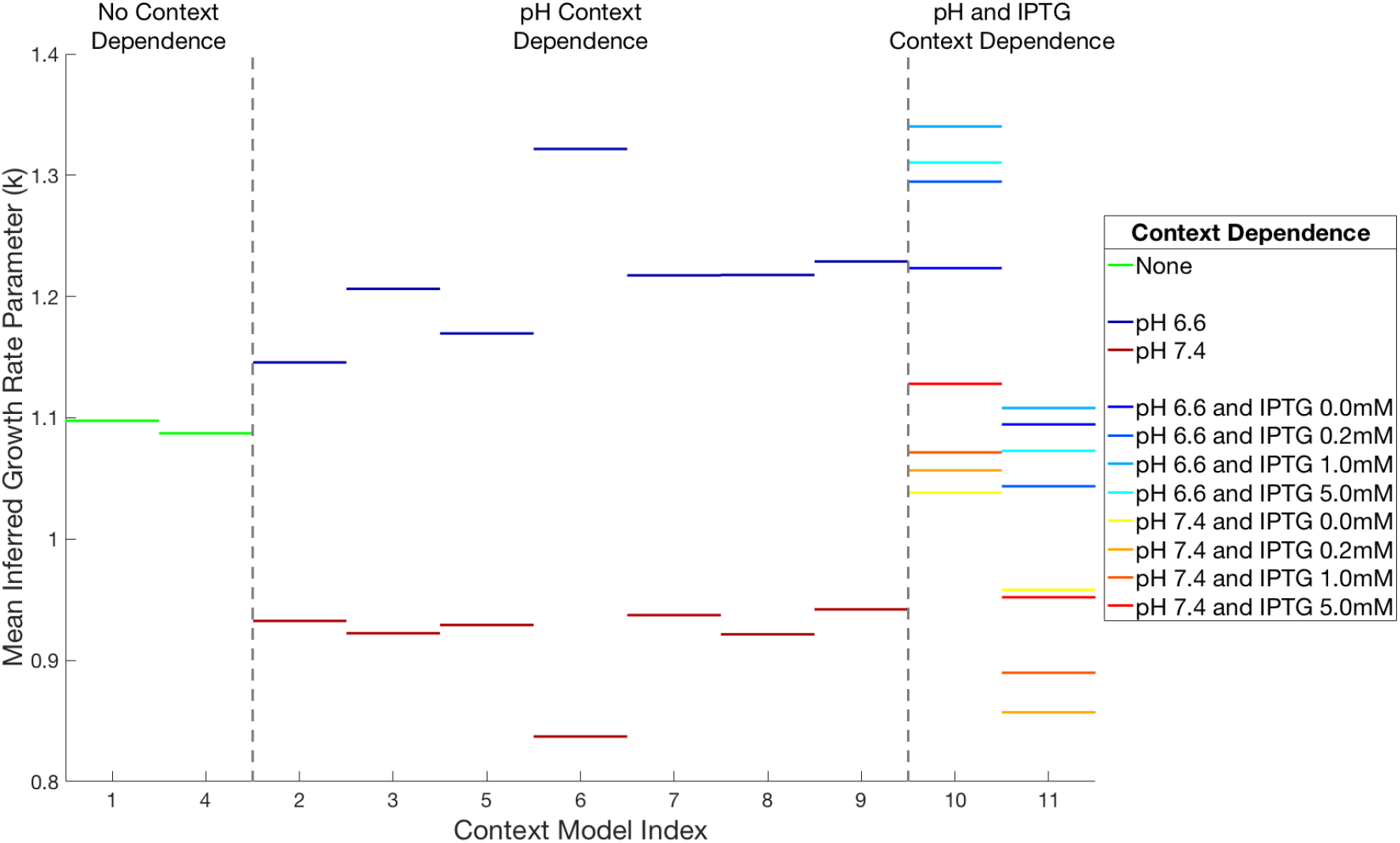
Comparison of the estimated mean growth rate for each context model in Table 3. We consistently observe that the higher pH value leads to an inferred lower cell growth rate across all models. We can also see that there is fairly significant variation in the inferred parameters, depending on context.

**Figure 8:**
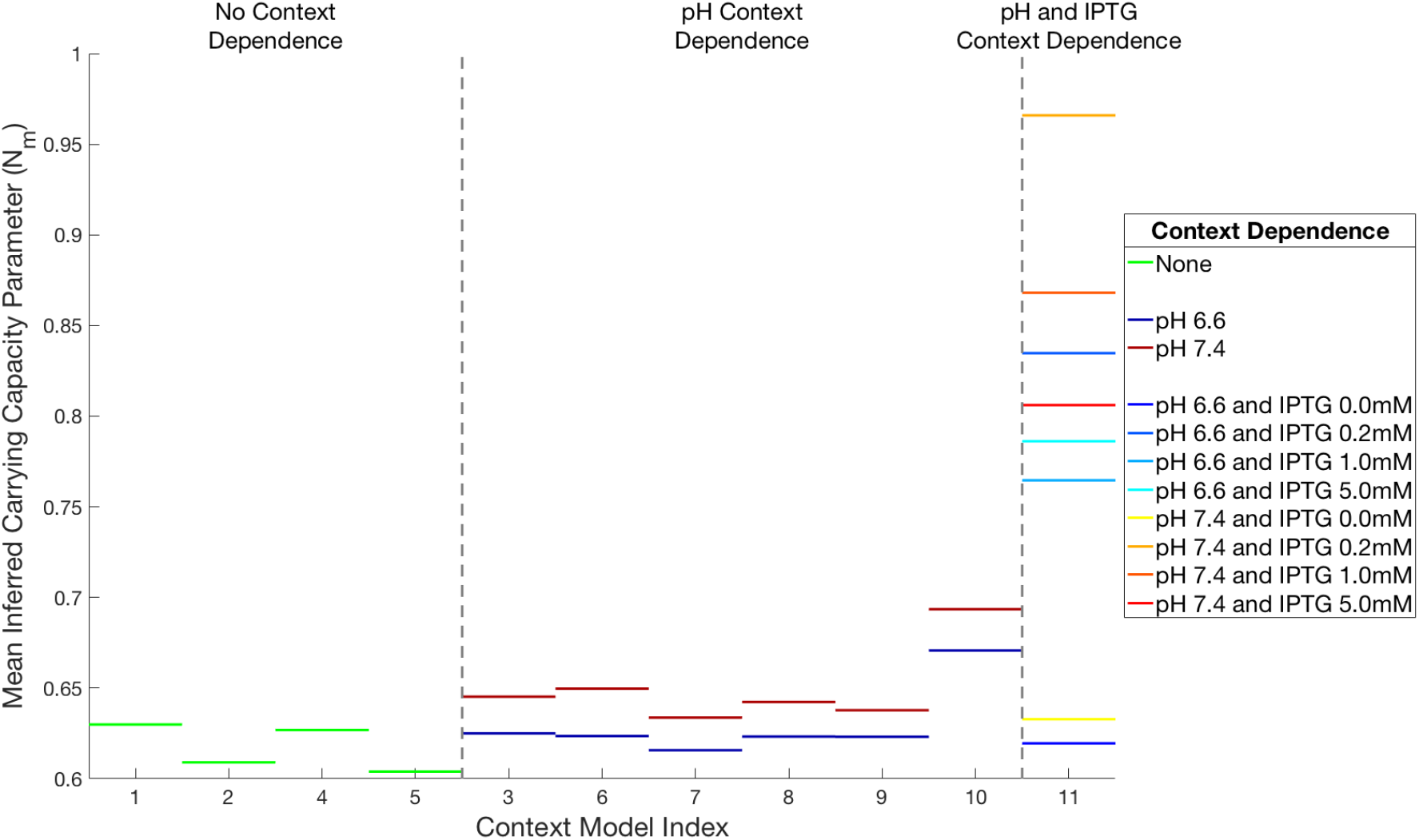
Comparison of the estimated mean carrying capacity for each context model in Table 3. With the exception of Model 11, we see that the higher pH value consistently leads to a higher inferred carrying capacity and that the estimated mean value is fairly consistent between the different context models. Models 10 and 11 show significant deviation from the other models, which can be due to over-fitting.

**Figure 9:**
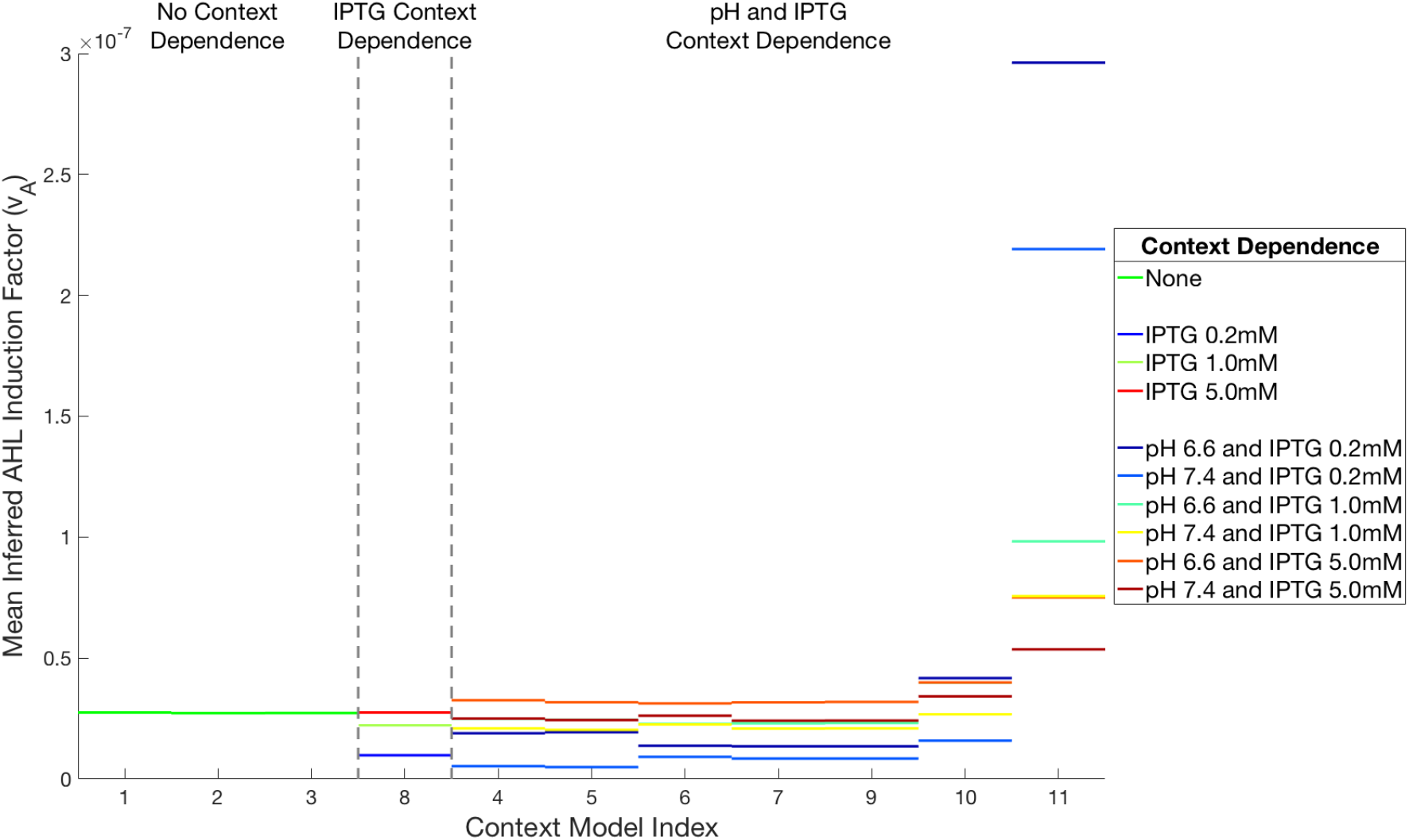
Comparison of the estimated mean AHL induction for each context model in Table 3. With the exception of Model 11, we find that the inferred AHL induction is fairly consistent across context models. We do observe a small dependence on IPTG value since lower levels of IPTG induction lead to lower AHL induction values.

After fitting the parameters for each context model, we compute the model’s plausibility (i.e. the Bayes factor). We compute the posterior relative probability of each model described in Table 3. We generally observe that there is significant context dependence since models with more context dependent parameters have much higher probability. Ultimately, the model with the most context dependent parameters effectively has all the posterior probability. Figure 15 shows the fit of the data using parameters from the posterior of the top three models, Models 9 - 11. We see that all these models capture the dynamics of the cell population well. However, we find that Model 11 can capture the dynamics in the data better than the other two models. The fitting error of each of the context models is illustrated in Figure 11.

**Figure 10:**
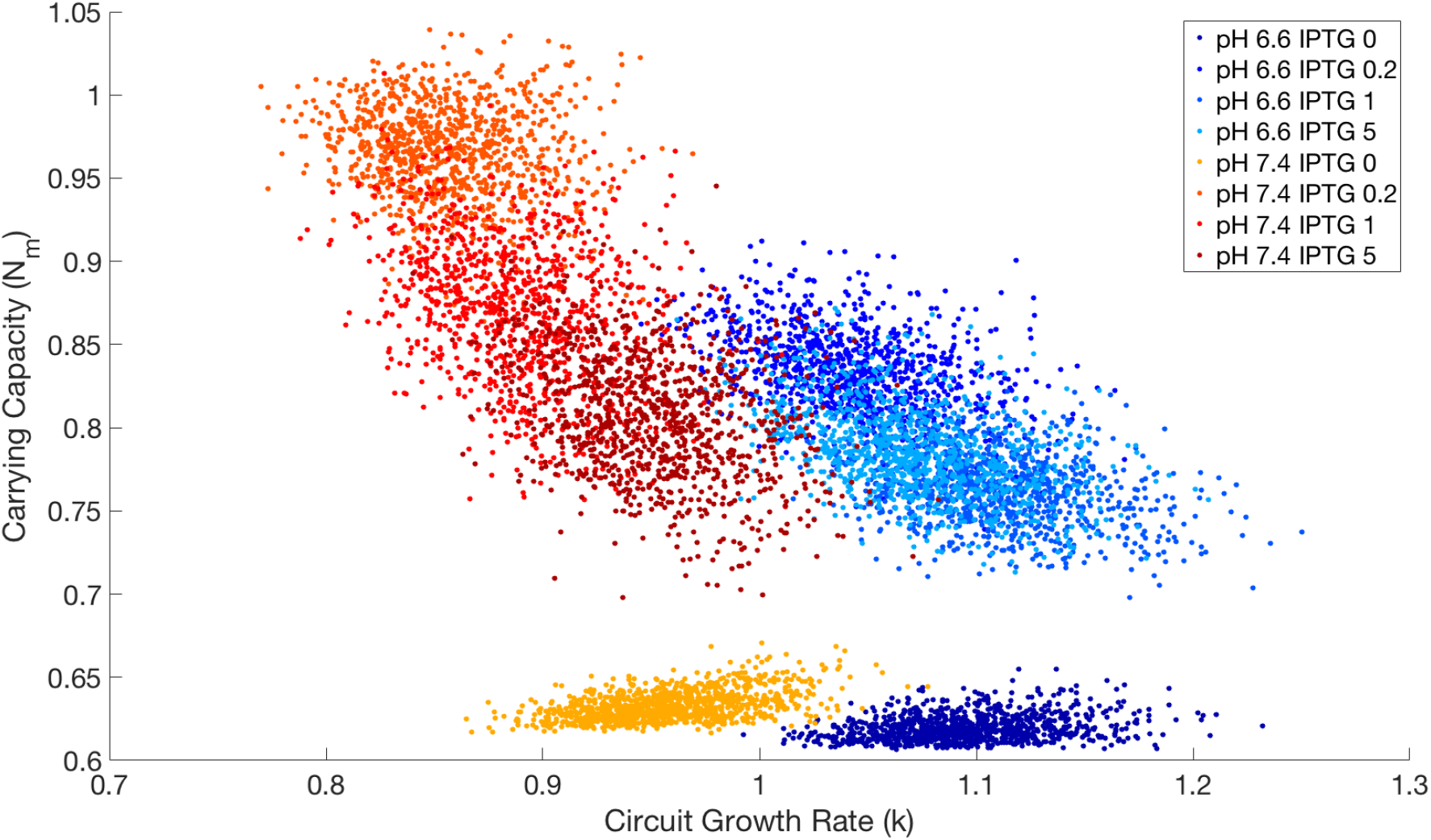
Posterior samples of the inferred growth rate and carrying capacity for experimental conditions with different IPTG induction levels and pH values. For Model 11 in Table 3, there appears to be significant context dependence on the pH and the IPTG values. The pH dependence appears as the separation between the cold colors (low pH) and the hot colors (high pH). It significantly affects the growth rate and less significantly affects the carrying capacity. Further, the individual colors show that there is also IPTG dependence, which is more apparent for the carrying capacity.

**Figure 11:**
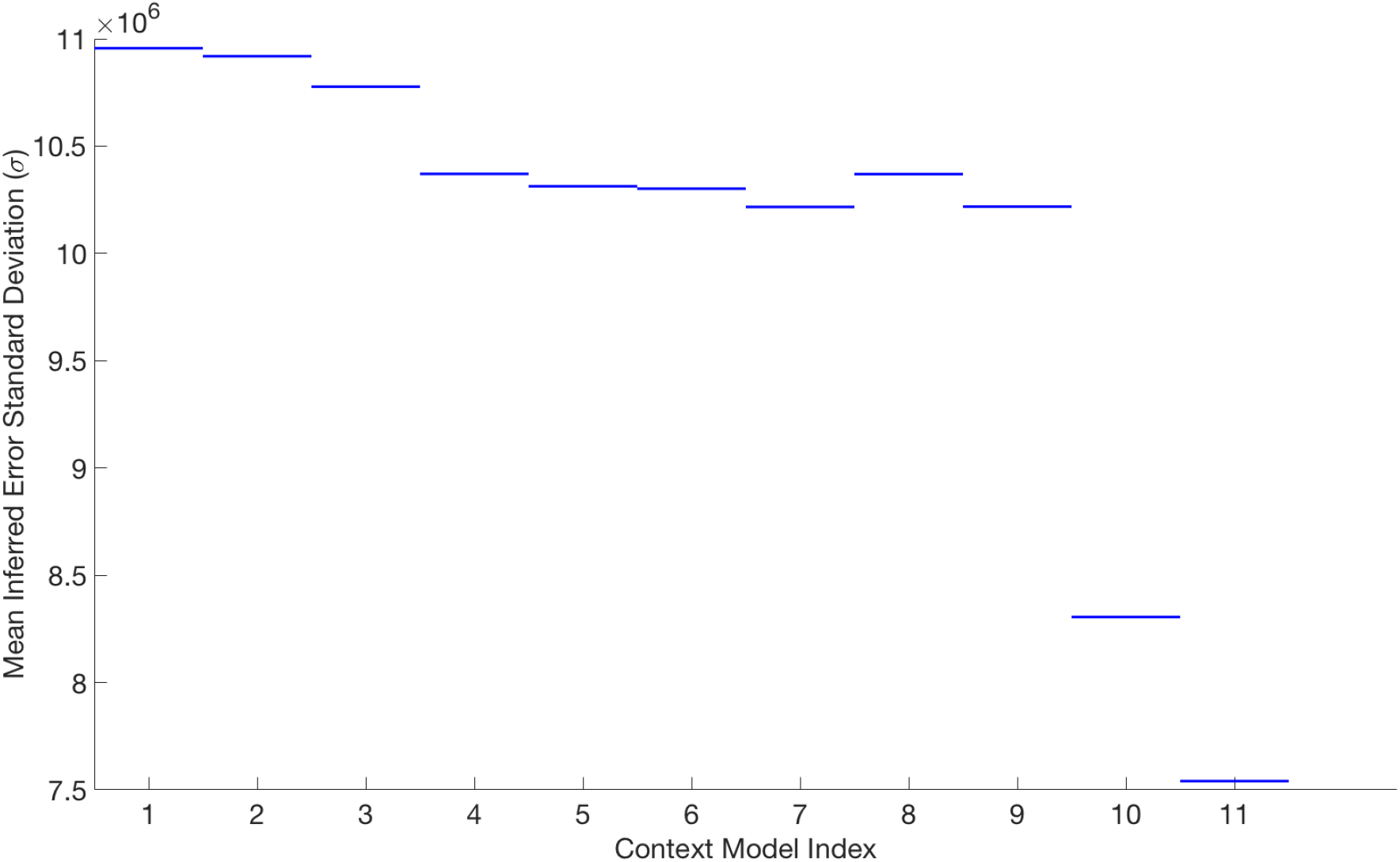
Comparison of the model fit error for each context model. In general, we see that increasing the complexity of the model by adding more context dependence leads to lower inferred error. The large reduction in error for Models 10 and 11 could indicate over-fitting since these two models add many new degrees of freedom.

In general, we see that more complex models can fit the data with smaller error. We also see significant reduction of the inferred error in Models 10 and 11. This could be evidence that we are over-fitting since the error levels are lower than what we could realistically expect our model to capture given the simplicity of the model and the complexity of the observed cell population response. While the Bayesian framework automatically incorporates a trade-off between the model complexity and the data fit quality, i.e. the Bayesian Occam’s razor, the conclusions we can draw from the Bayes factor analysis still require us to think critically about how reliable our models are. For instance, model selection is informing us that most of the parameters are highly context dependent. This could either mean that indeed they are dependent on both the pH and the IPTG values or that the growth circuit’s model in Equation (2) is not complex enough to capture the real dynamics. Hence, the added freedom we obtain by requiring more parameters to be context dependent helps us make up for deficiencies in the model of the cell growth circuit in Section 2.3. We believe this may be the case since data measurements of the growth circuit in stationary phase indicate that the cell population behaves unexpectedly, by either continuing to grow or to die; neither behaviors are captured by the cell growth circuit’s model. Ultimately, while we believe there is significant context dependence among some of the parameters, namely the growth rate, the carrying capacity, the AHL degradation rate, and the AHL induction, pinning down the exact dependence is difficult and will require additional targeted experiments built on what we have learned in this analysis.

## 5 Materials And Methods

Experiments generating the data were performed using the following materials and techniques.

### 5.1 Bacterial cell strains

*Eschericia coli* strain DH5*α*-Z1 was transformed with plasmid pLuxRI2, pluxCcdb3, or both to create two singly transformed control strains that contain each half of the circuit presented in You *et al*. and a doubly transformed strain that contains the necessary components to implement the circuit function. Plasmids pLuxRI2 and pluxCcdb3 were a gift of the Arnold laboratory [25] at the California Institute of Technology.

### 5.2 Cell growth experiments

Cell strains were inoculated into either LB or TBK (10g tryptone, 7g KCl per liter, 100mM MOPS to buffer) growth medium with kanamycin (50*µ*g/mL) and chlorampenicol (25*µ*g/mL) and allowed to outgrow overnight at 37°C. To begin the experiment, outgrown cells were diluted 200*×* into fresh media identical to their outgrowth media and 0.5mL of this suspension was pipetted in triplicate into each well of a square 96 well Matriplate (dotScientic, MGB096-1-1-LG-L).

Different concentrations of inducers were created by transferring the appropriate volume of a 500mM IPTG stock solution into the wells of a Matriplate using a Labcyte Echo 525 liquid handling robot. Volumes of IPTG were transfered to the plate *before* cell suspensions were added.

Plates with diluted cell suspensions were incubated at 37°C in a Biotek Synergy H2 incubator and plate reader with shaking for 21 and 28 hours, respectively, while absorbance at 600nm (OD600) was measured every 7 minutes. Cells were grown at 37°C with standard maximum linear shaking and lid and condensation gradient control.

### 5.3 Data processing

Due to different amounts of delay between when each of the cell populations has grown enough to begin being registered on the OD600 plate reader, the data was cut and aligned so that time zero corresponds to when the absorbance of a well exceeds a threshold. Data after a certain period is then also cut corresponding to when the cell population should be in stationary state. This is cut because the cell population behaves in ways not predicted by the model as time progresses in stationary state.

## 6 Conclusion

In this work, we investigated the context dependence of biological parameters in the You *et al*. cell growth regulation circuit under multiple experimental conditions. This context dependence captured the dependence of the biological parameters in the mathematical model of the circuit on the unmodeled context. We used Bayesian methods to estimate the model’s parameters and to compare different types of context-dependence relationships. We found that many of the parameters in the model of the growth control circuit exhibit significant context-dependence and that their estimated values changed significantly depending on pH levels and IPTG induction levels. Furthermore, we found that the estimated parameter values were significantly influenced by the assumed context-dependence. This highlighted the importance of considering context-dependence when performing parameter inference under an assumed context.

Developing better models and designing more informative experiments would help elucidate the effects of context dependence in the growth regulation circuit and would improve the applicability of Bayesian methods. First, the data used in this work is based on fluorescence measurements that cannot differentiate between living and dead cells. Both types of cells are counted, thus leading to bias in the cell population levels. This bias could be accounted for by using a mathematical model that tracks both living and dead cells and predicts the total florescence. Thus, we could closely track the true number of living cells for more accurate data measurements. Secondly, our results showed significant context dependence, but they were also affected by over-fitting dynamic behaviors in the cell population. The behavior of the cell population in stationary phase could not be captured by the current model of the circuit. We could either develop a more complex model of the growth regulation circuit that accounts for stationary phase behaviors or we could develop an uncertainty model of the data measurements that does not reward over-fitting. Finally, designing improved experiments could give us additional data to better infer parameters and context-dependence. Testing the system on a wider variety of pH levels, IPTG induction levels, different media, and different experimental setups could help us better isolate these effects.

## Acknowledgements

This project is partially sponsored by the Defense Advanced Research Projects Agency (Agreement HR0011-17-2-0008). The content of the information does not necessarily reflect the position or the policy of the Government, and no official endorsement should be inferred. This research was also funded through the John von Neumann Fellowship program at Sandia National Laboratories and used the computational resources of the National Energy Research Scientific Computing Center, a DOE Office of Science User Facility supported by the Office of Science of the U.S. Department of Energy under Contract No. DE-AC02-05CH11231. Sandia National Laboratories is a multimission laboratory managed and operated by National Technology & Engineering Solutions of Sandia, LLC, a wholly owned subsidiary of Honeywell International Inc., for the U.S. Department of Energy’s National Nuclear Security Administration under contract DE-NA0003525. SAND no. SAND2018-7077 J

## A Additional Data Measurements Of The Cell Growth Regulation Circuit

We compare mathematical models of the toxin-regulated and unregulated cell population circuits in [25] in Figure 12. When the circuit is not regulated by toxin-mediated cell death, the cell population quickly reaches carrying capacity. However, when the toxin regulates cell growth, the total cell population is kept below the carrying capacity. Thus, the circuit functions as a cell population cap.

**Figure 12:**
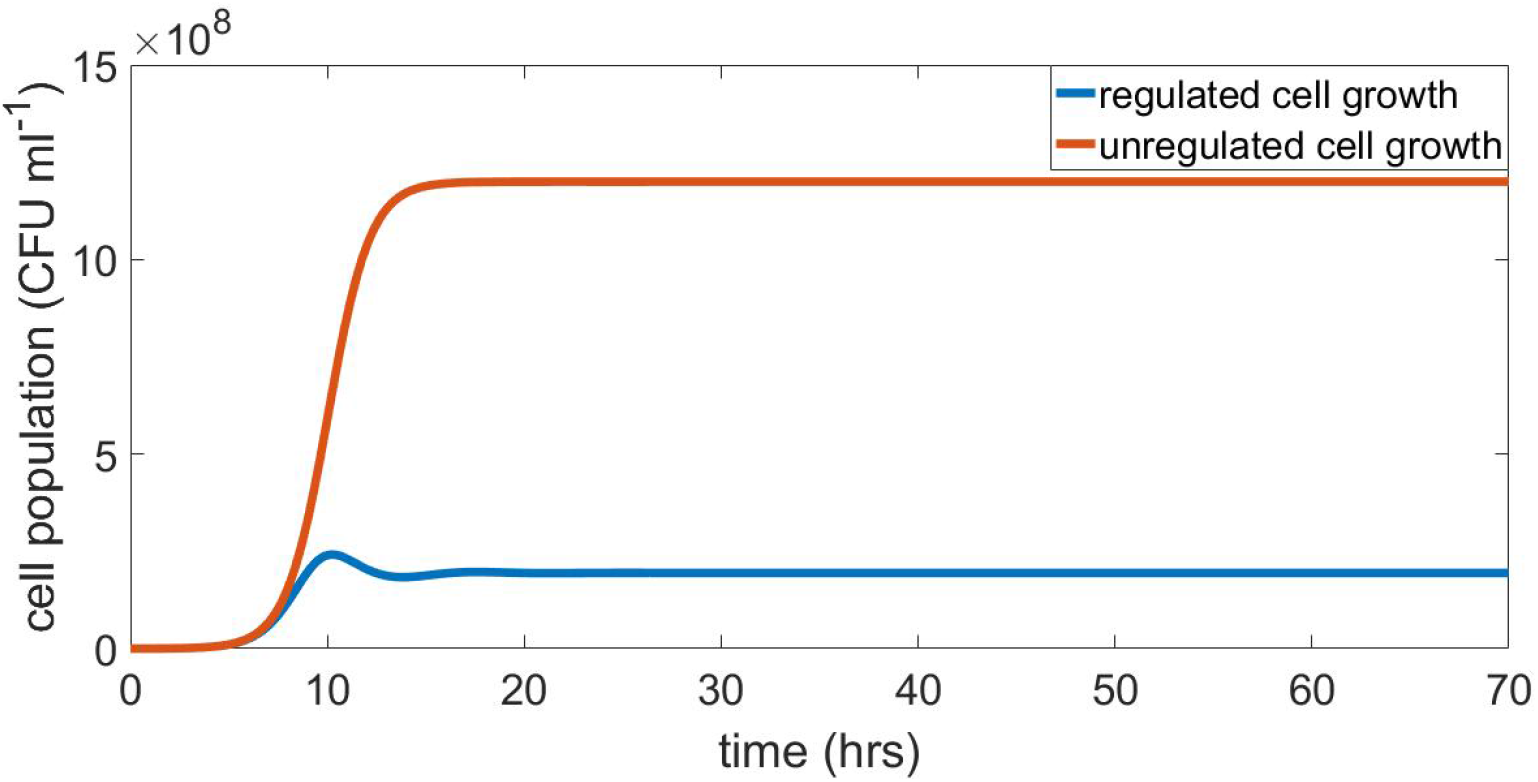
Comparing the unregulated cell population and the toxin regulated cell population. We compare mathematical models of the unregulated cell population and the toxin regulated cell population. The toxin CcdB sets the cell population cap lower than the unregulated cell population.

We plot the fluorescence data measurements of the growth regulation circuit at four induction levels in the TBK and the LB media in Figures 3 and 13, respectively.

**Figure 13:**
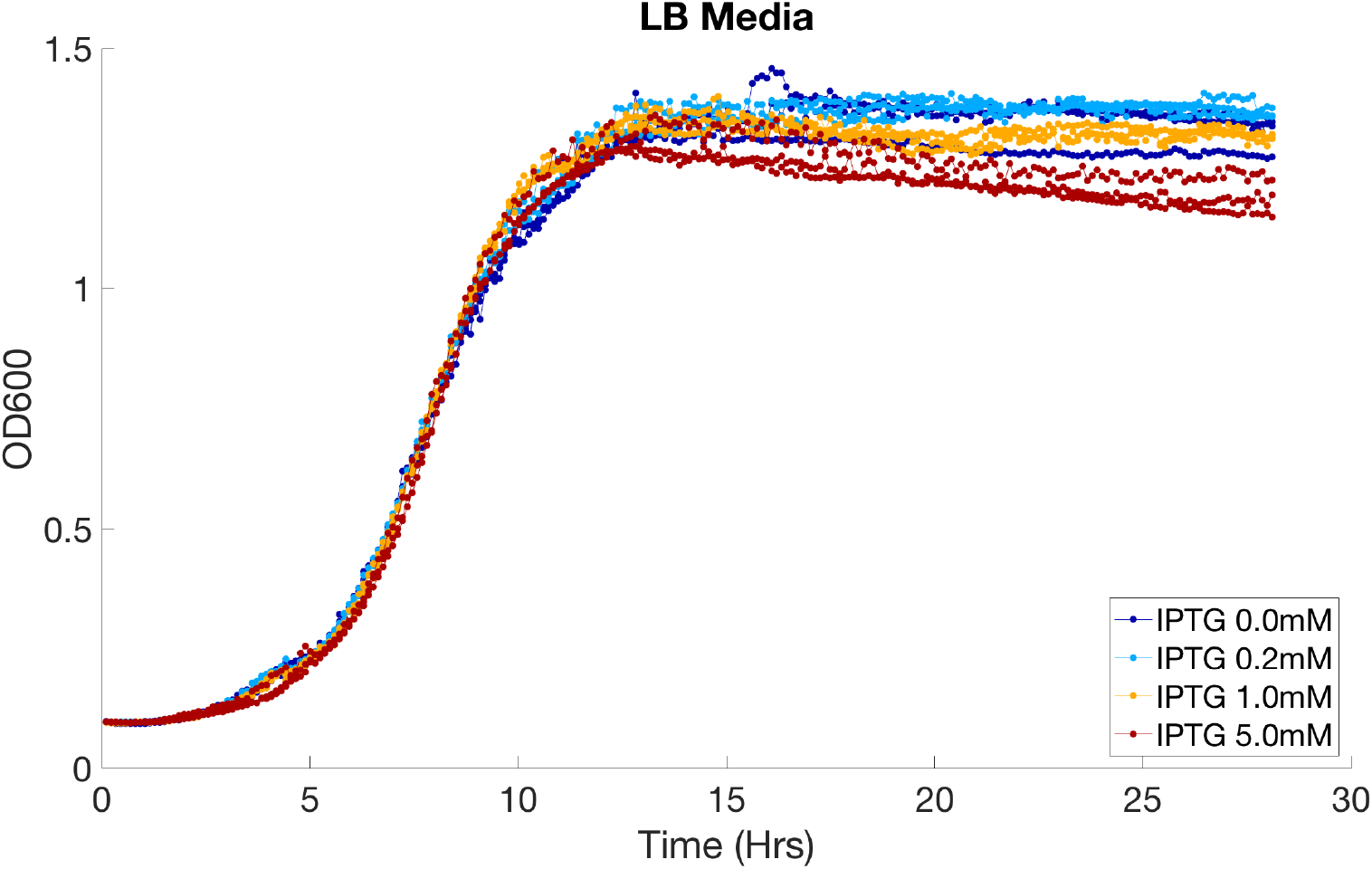
Fluorescence data measurements of the growth circuit in the LB medium. OD600 data measurements of the growth circuit with four different IPTG induction levels of 0mM, 0.2mM, 1.0mM, and 5.0mM in the LB medium.

The fits of the mathematical and context models found using Bayesian parameter inference for discussed in Section 4 for the control circuit experiments and IPTG and pH dependence experiments are presented in Figures 14 and 15, respectively.

**Figure 14:**
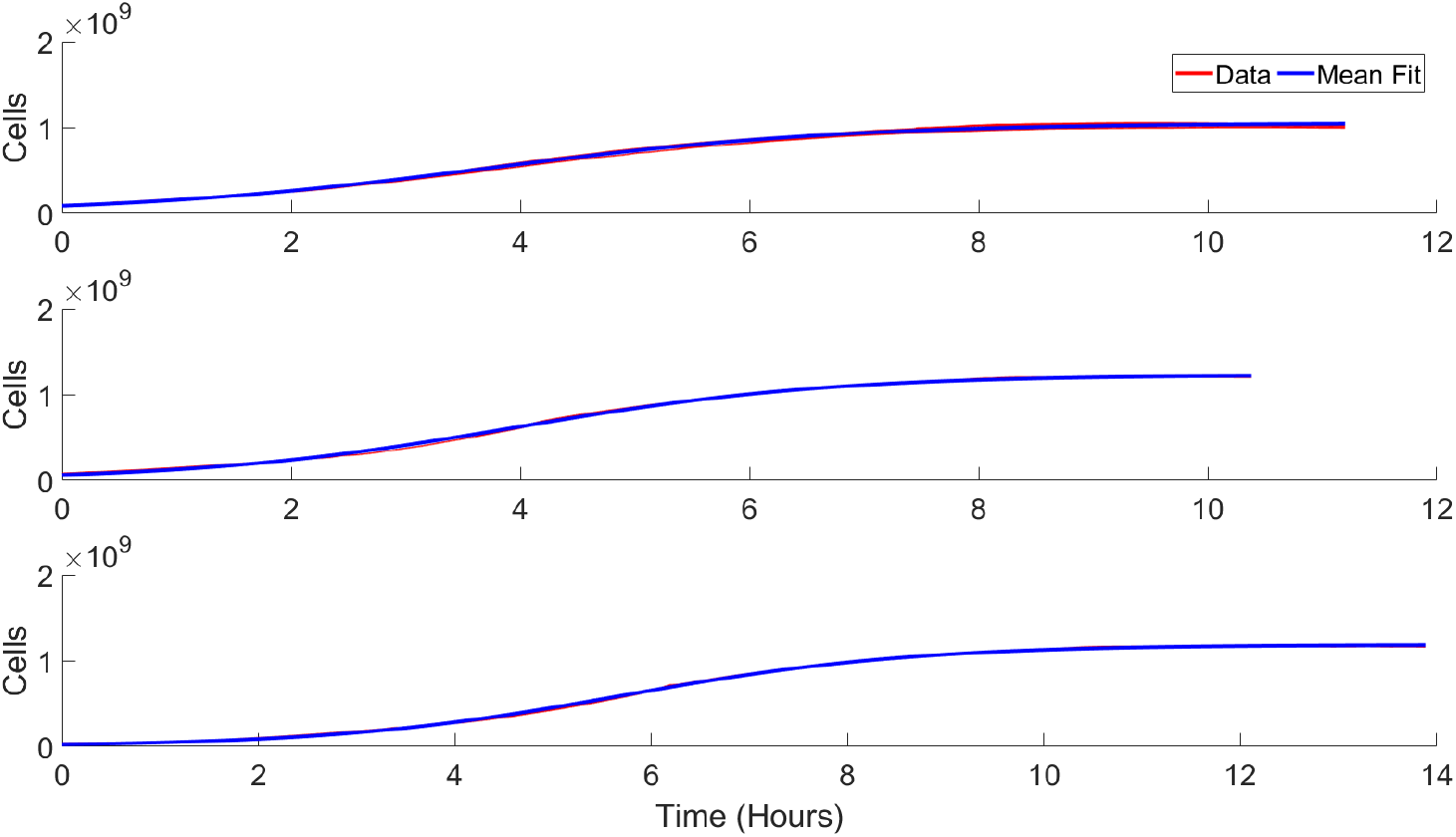
Comparison of the model fit to the measured data. For the control experiments, there is good agreement between the measured data and the mean fit to the data using the inferred parameters.

**Figure 15:**
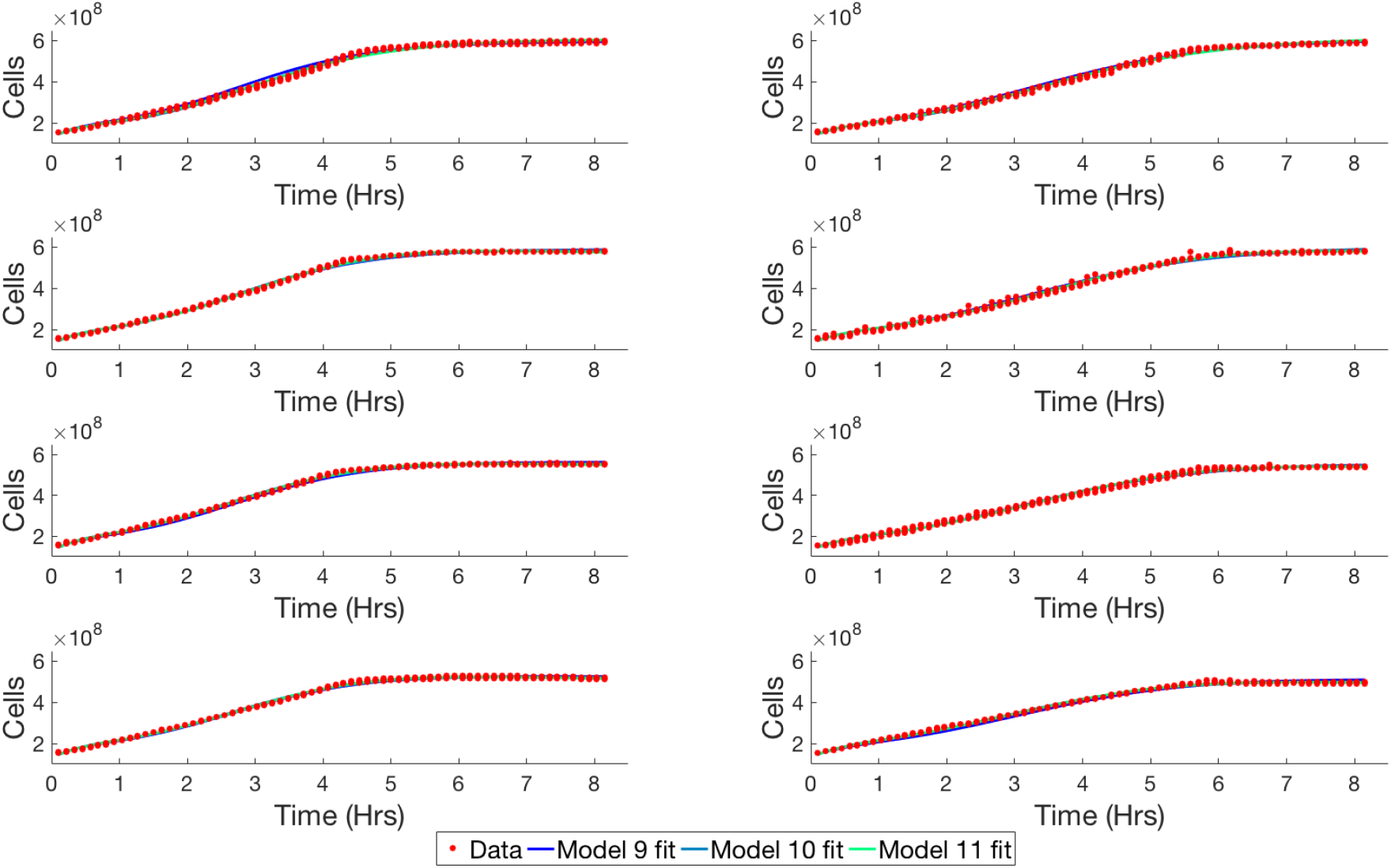
Comparison of the data fit with three different context models. When the growth rate, *k*, the carrying capacity, *N_m_*, and the AHL production factor, *v_A_*, are all pH and IPTG context dependent, we find that the fit of the model (green lines) to the data (red dots) can capture additional features that the other models are unable to, thus resulting in a high posterior likelihood of this context model.

## B Overview Of Bayesian Methods

### B.1 Prior Distributions

As discussed in Section 3, Bayesian parameter inference updates a probability distribution that describes model parameters using data. The initial probability distribution is called the prior and the updated probability distribution is call the posterior. The prior probability distribution captures our initial uncertainty about the model parameters, for example their range. When inferring both the mathematical model and the context model, we must describe prior distributions for both the biological parameters in the mathematical model and context parameters, known as hyperparameters, in the context model. The priors these parameters was chosen to reflect our knowledge given the experimental setup and reasonable ranges while in general trying not to be too informative.

Table 4 describes the prior distributions for the biological parameters in the mathematical model Equation (2). *k*, *N_m_*, *v_A_*, *d_A_*, *t*_0_, and *A*_0_ are properties of the dynamical system model. The priors for these parameters are defined with respect to hyperparameters discussed in Table 5. This is because these parameters may or may not exhibit context dependence and the hyperparameters are able to quantify the significance of the context dependence. The rest of the priors do not change with respect to the context model. *N* (0), *E* (0), and *A* (0) are initial conditions while *σ* is the model prediction error.

### B.2 Bayesian model selection

We describe how to perform Bayesian identification for the eleven model descriptions in Table 3. Bayesian identification for different mathematical model descriptions ℳ_*i*_ for *i* ∈ {1 … *N*} where *N* = 11, for our particular case, is similar to Bayesian parameter estimation, which was described in Section 3. We define a prior probability, *p* (ℳ_*i*_), for each possible model in the discrete set of models {ℳ_*i*_}_*i*=1…*N*_. Then we find the posterior probability of the model given the data, *p* (ℳ_*i*_ | 𝒟), by updating the prior according to the likelihood function of the data, *p* (𝒟 | ℳ_*i*_):

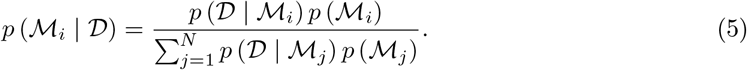

Unlike Bayesian parameter estimation, the normalization factor is easy to compute since it is a discrete sum over the unnormalized posterior probabilities for each model. However, the likelihood function for each model is computationally challenging because it requires solving the integral

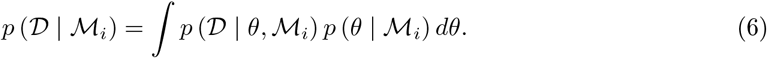

This integral is the normalization factor from Bayes’ theorem in Equation (4). This factor can be estimated using MCMC techniques discussed in Appendix B.4. The likelihood also enforces a natural trade-off known as the Bayesian Ockham’s razor in [3] as follows:

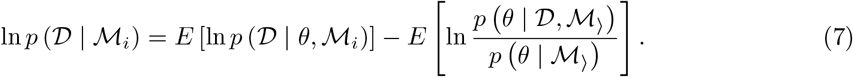

The first term captures the expected data fit, while the second term is the Kullback-Leibler divergence (i.e. information gain) between the prior and the posterior distributions. Therefore, the Bayesian Ockham’s razor trades-off how well the model fits the data with the information needed to be added to update the prior to the posterior.

### B.3 Bayesian model selection and hyperparameter models for identifying context dependence

As described in Section 3.2, we use probabilistic context models to capture the relationships between the biological circuit and its context. These probabilistic context models describe how mathematical parameters of the circuit model, *θ*, relate to one another and to the context of the circuit. The probabilistic context models are parameterized by hyperparameters, *ϕ*, that capture the biological context and the strength of the context relationship. Each context model ℳ_*i*_ for *i* ∈ {1 … *N*} has its own set of model parameters related by hyperparameters. Therefore, when estimating the Bayesian evidence for model selection between these different probabilistic relationship models, it takes the form of the following integral over the possible parameters and hyperparameters:

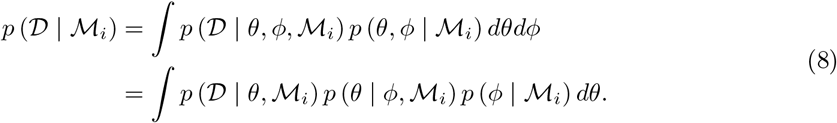

Further, to infer the context, we assume that the underlying structure of the mathematical model of the circuit does not change between context relationship models. Therefore, it must be that *p* (𝒟 | *θ*, ℳ_*i*_) = *p* (𝒟 | *θ*). Secondly, we assume that the prior on the hyperparameters, known as the hyperprior, is the same for each context model, i.e. *p* (*ϕ* | ℳ_*i*_) = *p* (*ϕ*). What changes between context models is *p* (*θ* | *ϕ*, ℳ_*i*_), the prior of the biological model parameters given the context model. With these assumptions, Equation (8) is equivalent to

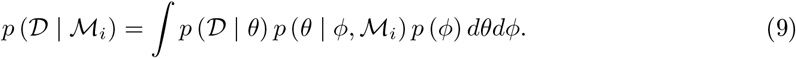

For example, in Figure 5, suppose the biological parameters *θ*_1_ and *θ*_2_ are dependent on the context in ℳ_1_, but independent of context in ℳ_2_. For ℳ_1_, the prior becomes *p* (*θ*_1_, *θ*_2_ | *ϕ*, ℳ_1_) = *p* (*θ*_1_ | *ϕ*) *p* (*θ*_2_ | *ϕ*) because context dependence implies that the biological parameters *θ*_1_ and *θ*_2_ are conditionally independent since they only depend on the context model parameters *ϕ*. For ℳ_2_, the prior becomes *p* (*θ*_1_, *θ*_2_ | *ϕ*, ℳ_2_) = *p* (*θ*_1_ | *ϕ*) 𝟙 (*θ*_1_ = *θ*_2_) = *p* (*θ* | *ϕ*). Because ℳ_2_ is context independent, we assume that the biological parameters are modeled using one parameter drawn from the hyperprior i.e. *θ* = *θ*_1_ = *θ*_2_ and therefore *θ*_1_ and *θ*_2_ are fully dependent as expressed by 𝟙 (*θ*_1_ = *θ*_2_) since their values do not change as the context changes.

Taking this general approach, for each of our context relationship models ℳ_*i*_, we construct the prior distribution *p* (*θ* | *ϕ*, ℳ_*i*_) and estimate the model evidence integral in Equation (9). Then we can compute the overall probability for each context model, as described in Equation (5).

### B.4 Markov Chain Monte Carlo methods

Markov Chain Monte Carlo (MCMC) methods are one of the primary ways of solving Bayesian parameter estimation and model selection problems. MCMC methods are necessary for sampling probability densities where the normalization constant, *p* (𝒟) in Bayesian inference, is very difficult to compute. Using MCMC methods, we construct a Markov chain whose stationary distribution is the posterior distribution *p* (*θ* | 𝒟) that we are interested in sampling. Using samples from this population, we can then estimate properties of the distribution and compute expectations. Unlike Monte Carlo sampling, these samples are correlated since they are coming from a Markov process. However, by appropriately designing the sampler, we can minimize the correlation and understand its impact on estimates using the Markov chain central limit theorem.

The most common form of MCMC is the Metropolis-Hastings algorithm, which creates a reversible Markov chain with the desired stationary distribution. This is achieved using a proposal distribution *Q* (*θ*′ | *θ*) for proposing new samples and an acceptance probability *α* (*θ*′ | *θ*) for deciding to accept the proposed samples:

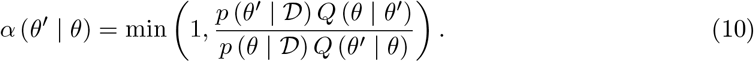

The key insight is that this acceptance rule only relies on the ratio of the probability densities *p* (*θ*′ | 𝒟) and *p* (*θ* | 𝒟), so the normalization constant is not needed. For more details on the Metropolis-Hastings sampler and MCMC methods in general, see [4].

Solving the Bayesian model selection problem requires a more sophisticated implementation of MCMC because we need to estimate the normalization constant *p* (𝒟 | ℳ). The key is to view Equation (6) as an expectation as follows:

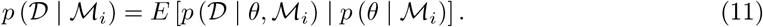

Hence, we must estimate the expected value of the likelihood given samples from the prior. This can be achieved through the combination of Sequential Importance Sampling and MCMC [6, 8]. We employ the Sequential Tempered MCMC algorithm described in [8, 9].

## C Posterior biological parameter histograms for different context models

Figures 16 – 18 provide posterior histograms for the inferred growth rate, carrying capacity, and AHL induction factor under different assumed context models. We observe that depending on the context model, the inferred parameter values may be significantly different to the extent that the distributions may not overlap at all. This is the case for the growth rate parameter and it highlights the need for considering context models when estimating the values of biological parameters.

**Figure 16:**
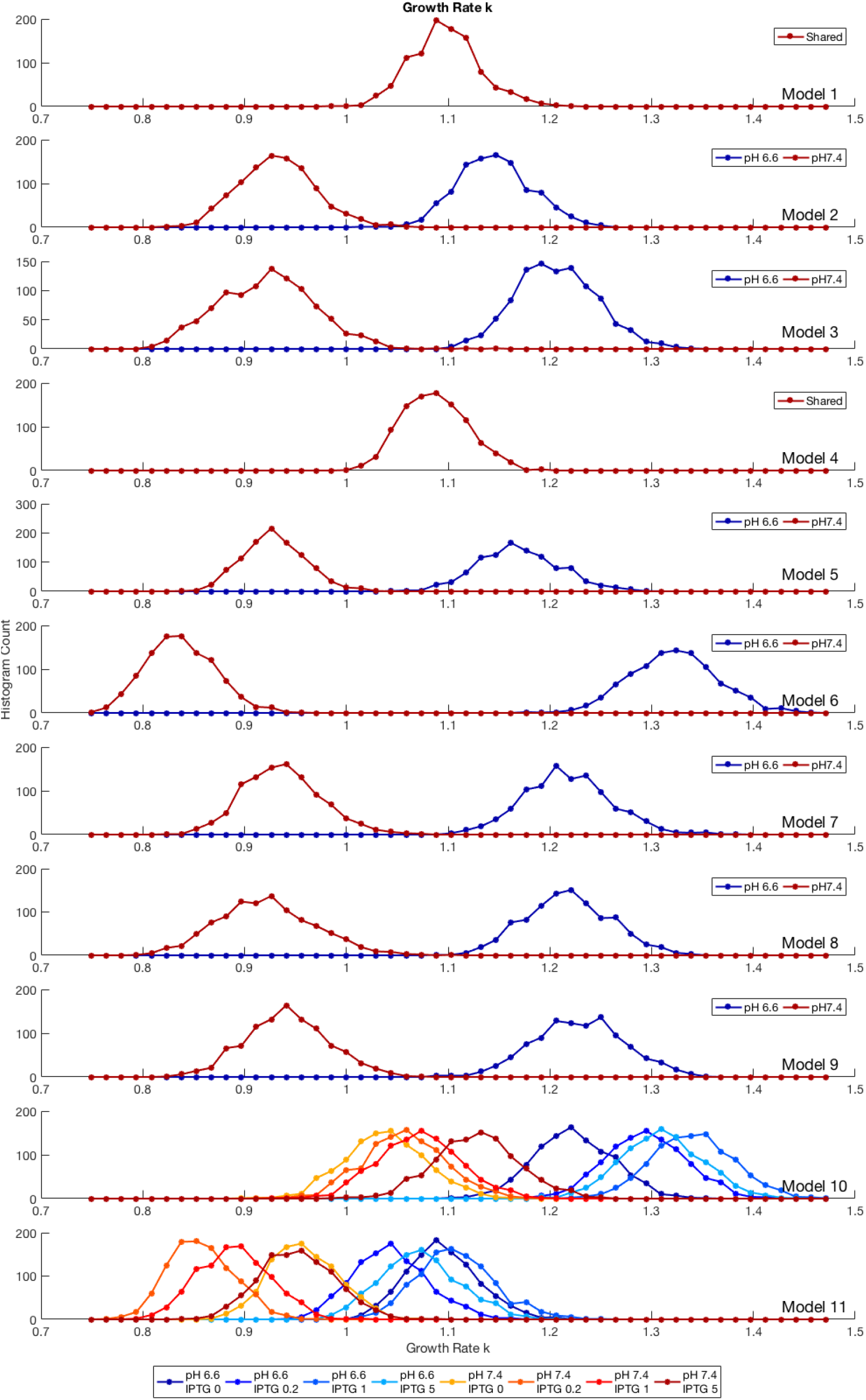
Histograms of the inferred growth rates for different context models. Depending on the context model, the growth rate is either independent of context (1 and 4), dependent on the pH value (2, 3, 5, 6 *−* 9), or dependent on both the pH and IPTG induction values (10 and 11). For each model, colors represent the posterior histogram of growth rate parameter values for the different experimental conditions.

**Figure 17:**
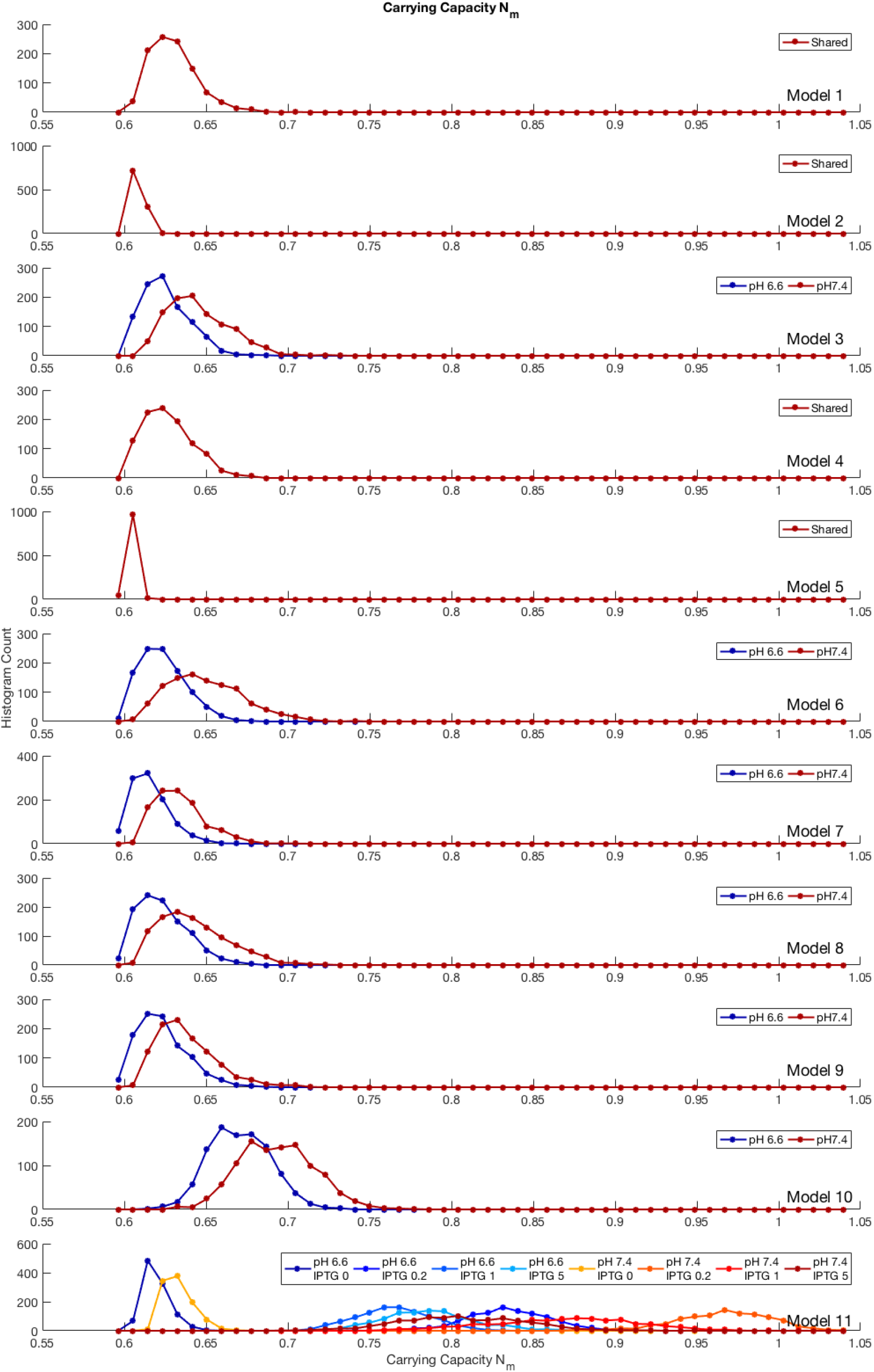
Histograms of the inferred carrying capacity for different context models. Depending on the context model the carrying capacity is either independent of context (1, 2, 4, and 5), dependent on the pH value (3, 5, 7 *−* 10), or dependent on both the pH and IPTG induction values (11). For each model, colors represent the posterior histogram of carrying capacity parameter values for the different experimental conditions.

**Figure 18:**
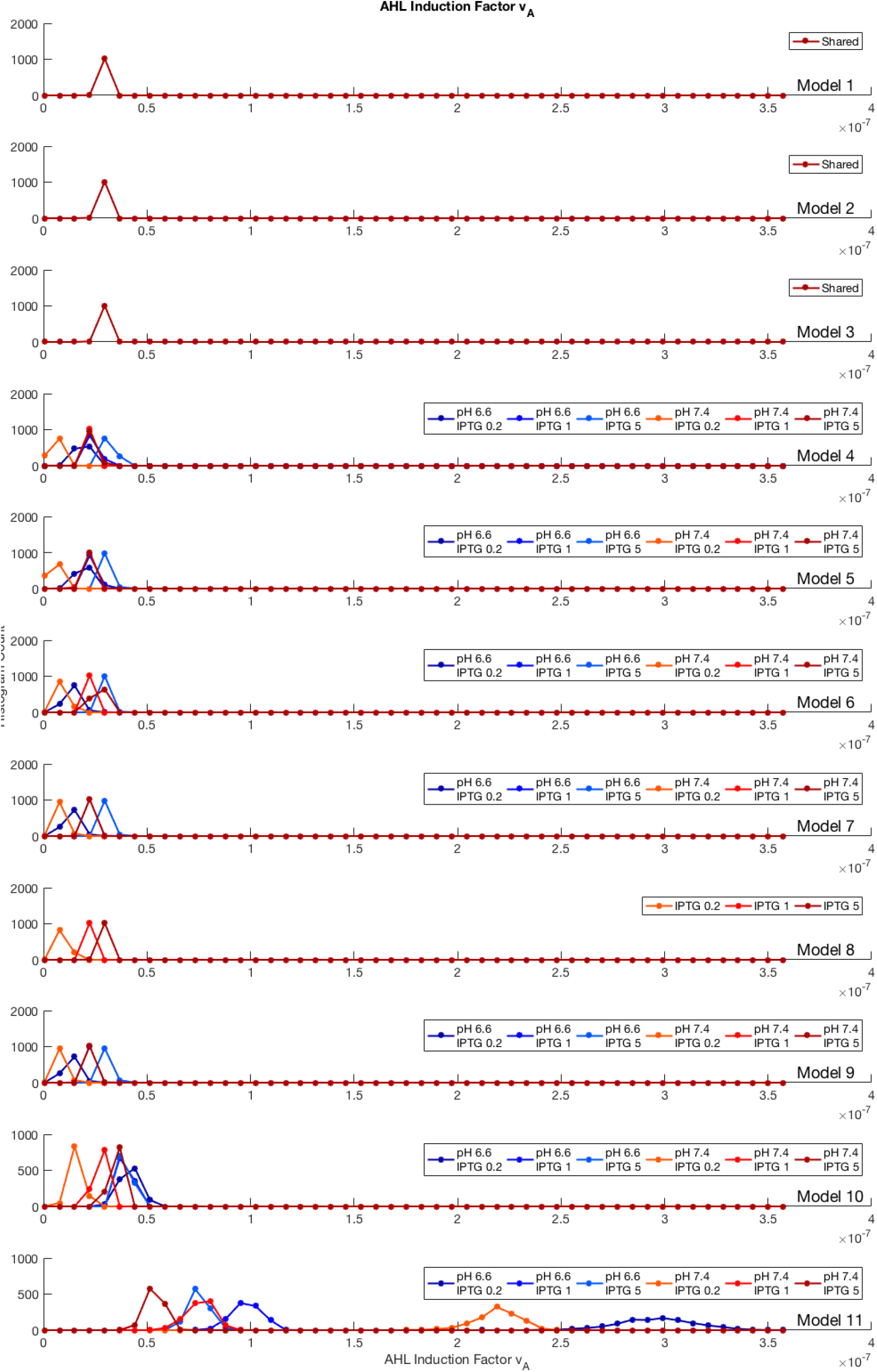
Histograms of the inferred AHL induction for different context models. Depending on the context model, the AHL induction is either independent of context (1 *−* 3), dependent on the IPTG induction value (8), or dependent on both the pH and IPTG induction values (4 *−* 7, 9 *−* 11). For each model, colors represent the posterior histogram of AHL induction parameter values for the different experimental conditions.

